# A Mixed T2/T17-Associated Systemic Immune Signature Links Airborne Pollutant Exposure to Persistent Respiratory Symptoms

**DOI:** 10.64898/2026.08.17.745275

**Authors:** Armando M. Marrufo, Christine H. Wendt, Eric Garshick, Vincent S. Fan, Raúl San José Estépar, Li-Zhen Song, Jiayi Li, Sarvika Periyapalayam Murali, Ivan M. Marrufo, Maxwell Stewart, Demerise Johnston, David B. Corry, Tianshi David Wu, Farrah Kheradmand

## Abstract

**Background:** The systemic immune responses associated with persistent respiratory symptoms (PRS) after exposure to airborne environmental pollutants remain poorly understood.

**Objective:** To identify immune disturbances associated with PRS, defined as persistent wheeze, cough, or breathlessness, we examined systemic immune responses and airway function in a cross-sectional cohort with detailed histories of airborne pollutant exposure.

**Methods:** Never-smoking post-deployment Veterans with PRS (n=16) or without PRS (n=24) underwent chest computed tomography, pulmonary function testing, and oscillometry to assess structural and functional airway abnormalities. Peripheral blood mononuclear cells (PBMCs) were stimulated with anti-CD3/CD28 antibodies, lipopolysaccharide, or β-glucan, and cytokine production was measured. Correlation analyses evaluated associations between cytokine responses and physiological measures of airway function.

**Results:** Oscillometry, but not conventional pulmonary function testing or chest computed tomography, detected small-airway abnormalities in participants with PRS, including significantly greater frequency dependence of resistance and higher resonant frequency. Baseline PBMC cytokine concentrations were similar between groups. After stimulation, however, PBMCs from participants with PRS showed increased IL-17A production consistent with a type 17 (T17) response; innate stimulation also increased the type 2 (T2) cytokines IL-33 and IL-4. T2/T17 cytokine responses correlated positively with oscillometric measures of small-airway dysfunction.

**Conclusion:** Individuals with PRS exhibited a stimulus-dependent systemic T2/T17 immune signature that was associated with early small-airway dysfunction.

**Clinical Implication:** Stimulus-dependent systemic immune profiling, combined with oscillometry, may help identify early respiratory abnormalities in pollutant-exposed individuals whose conventional pulmonary tests remain normal.

**KEY MESSAGES:**

- Oscillometry detected early small-airway dysfunction in pollutant-exposed individuals with persistent respiratory symptoms, whereas conventional pulmonary function tests and chest CT did not distinguish the groups.
- Immune stimulation revealed a mixed T2/T17 systemic signature in participants with persistent respiratory symptoms, including increased IL-17A and stimulus-dependent increases in IL-4 and IL-33.
- T2/T17 cytokine responses correlated with oscillometric abnormalities, linking systemic immune dysregulation to small-airway dysfunction and suggesting a potential approach for identifying early pollutant-associated respiratory disease.

**CAPSULE SUMMARY:** A stimulus-dependent systemic T2/T17 immune signature correlated with oscillometric evidence of small-airway dysfunction, identifying a potential early respiratory phenotype in pollutant-exposed individuals with persistent respiratory symptoms.

## INTRODUCTION

Long-term inhalation of airborne pollutants, originating from both occupational and environmental sources, has been established as a contributing factor to the development of various respiratory symptoms, including chronic cough, wheezing, dyspnea, and increased sputum production (1–4). However, less is known about risk factors associated with disease progression (5), highlighting a significant unmet need in factors that distinguish early lung disease development associated with air pollution, particularly within populations considered at risk. Many persistent lung diseases, including chronic obstructive pulmonary disease (COPD), asthma, interstitial lung diseases (ILDs), and bronchiectasis, are associated with smoking, poor air quality, and individual susceptibility factors, causing increased morbidity and mortality globally (6, 7). Notably, air pollutants have the capability to traverse the alveolar-capillary barrier, subsequently modulating the immune system and promoting disease development (8, 9). The relationship between occupational and environmental exposures and clinical outcomes highlights the central role of the lungs in influencing systemic immunity (10, 11).

The innate and adaptive immune systems are critical in maintaining lung homeostasis and orchestrating the inflammatory and reparative processes necessary for respiratory health. Within these systems, specific T helper (Th) cell subsets and innate immune cells contribute distinct functions. Type 1 (T1) immunity, which includes Th1 cells and innate immune cells such as macrophages and neutrophils, produce interferon-gamma (IFN-γ), which is essential for defending against intracellular pathogens (12). In contrast, type 2 (T2) immune responses, include Th2 and innate lymphoid cells (ILCs), secrete canonical cytokines, such as IL-4, IL-5, IL-9, and IL-13. These cytokines help resolve inflammation and facilitate tissue repair (12). Type 17 (T17) immunity, marked by the presence of Th17 cells, expressing IL-17A, is associated with antimicrobial defense against extracellular pathogens, like bacteria and fungi, as well as chronic inflammation (12–17). Persistent immune responses to environmental stimuli can disrupt normal immune regulation, resulting in severe lung and systemic inflammation, autoimmune disorders, or chronic infections (18). Furthermore, exposure to air pollutants can impact not only the lungs but also the vasculature and other organs, suggesting that occupational and environmental exposure may provoke immune responses extending beyond the respiratory system (19–21). Despite these insights, the precise role of these exposures in modifying innate and adaptive immune responses among at-risk populations remains to be fully understood.

Military personnel deployed to land-based environments serve as a well-established example of individuals exposed to a broad range of inhalational pollutants, illustrating the long-term effects of persistent air pollution on lung health (22). Many veterans experience persistent respiratory symptoms following these deployments including chronic wheezing, cough, and breathlessness (23). To better understand how deployment-related airborne exposures impact systemic immunity, we analyzed data from Veterans participating in the Veterans Affairs (VA) Cooperative Studies Program #595 and ancillary <u>L</u>ung <u>E</u>ffects of <u>D</u>eployment <u>E</u>xposure (LEDE) study (2, 24). This approach allowed us to examine immune responses and lung function in Veterans with and without persistent respiratory symptoms. Through this model, we identified key associations between altered adaptive and innate systemic immune responses and respiratory symptoms. These findings, along with physiological measurements, differentiated Veterans with persistent symptoms from those without, providing valuable insight into the effects of chronic air pollution exposure on systemic immunity and lung function.

## METHODS

### Study population and recruitment

Participants were randomly selected for the parent Service and Health Among Deployed Veterans (SHADE; NCT02825654, VA Cooperative Studies Program #595: Pulmonary Health and Deployment to Southwest Asia and Afghanistan) study using the Department of Defense (Defense Manpower Data Center deployment roster records), as part of meeting the inclusion criteria for the SHADE studies (2). Briefly, the SHADE study to assess respiratory health and cumulative inhalational exposures in Veterans deployed to land-based military service between October 2001 to August 2021 (2). This study specifically assessed lung function, prevalence of persistent respiratory symptoms, asthma, sinusitis, and rhinitis in this observational cohort. All SHADE participants underwent an in-person interview, which consisted of completing a structured questionnaire, including demographics, smoking history, health, military service-related information (service branch, occupation, location, and duration of deployments), and deployment-related exposure assessment.

The Lung Effects of Deployment Exposure (LEDE) study, a subset of the SHADE study, specifically investigates how specific pollutant exposures from burn pits and other environmental and occupational sources affect systemic biomarkers, lung structure, and physiological measurements. To minimize the effects of cigarette smoking, recruited participants from the SHADE study with a history of smoking (cigarettes, pipes, or vapes) were excluded from the LEDE study. The LEDE participants with persistent respiratory symptoms, are defined as those with: (a) chronic cough, (b) chest wheezing or whistling in the previous 12 months, or (c) dyspnea (breathlessness not due to strenuous exercise). LEDE participants who displayed at least one or more of these persistent symptoms described were selected as the part of the symptomatic group. Participants who did not display any of these three persistent respiratory symptoms were selected as part of the asymptomatic group. For the study presented here, we chose 40 never-smoker participants from the LEDE study performed at four study sites (Boston, Houston, Minneapolis, and Seattle VA medical centers) who displayed distinct clinical presentations, without (n=24, asymptomatic group) and with persistent respiratory symptoms (n=16), for analyzing immune response and lung physiological measurements (e.g., spirometry, oscillometry, and lung CT quantification). Together, data collected from the SHADE and LEDE studies include a respiratory health questionnaire, information about smoking history, military and civilian occupational exposures, jobs, medical history, demographics (age, sex, and race), PFTs, oscillometry, quantitative CT images, white blood cell (WBC) differential count, and cytokine production levels isolated from Veterans’ PBMCs.

### Exposure and respiratory symptom assessment

A comprehensive exposure assessment consisting of a 32-item questionnaire regarding occupational and environmental deployment-associated exposures was used (2). The exposure questionnaire focused on repeated pollutant exposures to military occupation-related vapors, gas, dust, or fumes (VGDFs); burn pit smoke, including regular exercise and work at or near burn pits; open air combustion particulates, including exposures to military activities, oil well, and refinery fires; and other toxicants, such as pesticides and chemical warfare agents (2). Heavy exposure in the questionnaire is defined as direct or prolonged exposure or could clearly be sensed at the time (e.g., effects felt on throat, eyes, or breathing). Those with affirmative answers were asked for duration by total number of months and days per month that exposure occurred.

The health component of the interview consisted of symptom assessment, based on the American Thoracic Society (ATS) and Division of Lung Diseases, National Heart, Lung, and Blood Institute (ATS-DLD-78) respiratory and National Health and Nutrition Examination Survey (NHANES) (25). All studies were approved by the Institutional Review Boards of all participating VA centers and institutions, and written consent was obtained from all study participants.

### Pulmonary Function Tests

All PFTs were performed at the participants’ VA medical center study sites using standard equipment. Spirometry was performed following both ATS and European Respiratory Society (ERS) guidelines by trained staff (26–28). PFT measurements included percent predicted of forced expiratory volume in one second (FEV_1_), forced volume capacity (FVC), and FEV_1_/FVC ratio, which were acquired before and 15-30 minutes after two doses of bronchodilator (albuterol, 180 µg total). Calculation of all PFTs were normalized to reference values based on ATS/ERS guidelines. The lung diffusion test was utilized to measure diffusion capacity for carbon monoxide (DL_CO_). Lung volumes (total lung capacity, TLC) were assessed by multiple breath nitrogen washouts. All values for each parameter were determined and normalized using the reference equations based on age, height, race, and birth sex (29).

### Measurement of Small Airways’ Function using Impulse Oscillometry

Oscillometry measurements were performed using impulse oscillometry (IOS) standardized protocol (30). Six standard variables measured as % predicted pre-bronchodilator use included: total airway resistance at 5 Hz (*R_5_*), central airway resistance at 19 Hz (*R_19_*), reactance at 5 Hz (*X_5_*), resonant frequency (*F_res_*), and reactance area (*AX*). The frequency dependence of resistance (*R_5_-R_19_*) was reported as cm H_2_O/(L/s).

### Quantitative Computed Tomography Morphology of Lungs

Lung structure was measured on CT scans of the lungs obtained from each subject in the supine position. Percent emphysema (% low attenuation area [LAA]), gas trapping, and airway wall thickness (Pi10, internal perimeter of 10 mm) were determined from imaging using Parametric Response Map, a voxel-wise image analysis technique (31). CT acquisition was performed at end-inspiration and normal-expiration for each subject using standardized published protocols (31, 32). Quantitative analysis was based on reconstructed data using the standard reconstruction kernel. Quantitative analysis of emphysema was assessed on segmented lung parenchyma images using Slicer measured as Hounsfield units. Emphysema was assessed by segmenting the lung parenchyma from the chest wall and central blood vessels as described above. Airway thickness represents the mean airway wall thickness normalized to a ‘theoretical’ airway lumen of 10-mm inner perimeter (33–36).

### White Blood Cell Differential Count and Peripheral Blood Mononuclear Cell Isolation

Peripheral blood samples were collected by standard venipuncture into sterile tubes containing ethylenediaminetetraacetic acid (EDTA). Differential percent and absolute WBC counts, monocytes, neutrophils, eosinophils, basophils, and lymphocytes were obtained from clinical laboratories.

Peripheral blood samples were obtained from subjects at the study site in gradient-separated and whole blood and processed immediately upon receipt for isolation and cryopreservation of PBMCs. The plasma was aspirated without disturbing the cellular interface, transferred into sterile conical tubes, and diluted with 1X phosphate-buffered saline (PBS). Samples were centrifuged at 1500g for 5 minutes at 4°C to obtain the cell pellets. To remove residual erythrocytes, the cell pellet was resuspended in sterile ammonium-chloride-potassium (ACK) red blood cell lysis buffer (0.15 M NH_4_Cl, 10 mM KHCO_3_, 0.1 mM Sodium EDTA, and distilled water, pH adjusted to 7.2-7.4 with 1N HCl) and incubated for 3 minutes at room temperature. A small aliquot of the cell suspension was diluted 1:10 with trypan blue solution (Corning) for viable cell counting using a hemocytometer. Following incubation, Roswell Park Memorial Institute (RPMI) 1640 cell culture media supplemented with 2.05 mM L-glutamine (Cytiva), 10% fetal bovine serum (FBS, ThermoFisher), and 1X penicillin/streptomycin (Corning) was added to neutralize the effect of lysis buffer. The suspension was centrifuged at 1500g for 5 minutes at 4°C, and the cell pellet was resuspended in freezing medium consisting of 80% FBS and 10% dimethyl sulfoxide (DMSO, Sigma Aldrich) prepared in complete medium. The entire suspension was transferred into sterile cryovials and stored at -80°C overnight to allow gradual freezing and stored in liquid nitrogen for batch testing.

### Ex Vivo Stimulation of PBMCs

PBMCs were thawed in batches, washed with sterile 1X PBS without calcium and magnesium (Corning) three times before final resuspension in supplemented RPMI 1640 cell culture media (Cytiva). A small volume of the cell suspension was stained with trypan blue solution for viable cell counting using a hemocytometer. Viable PBMCs from each subject in the asymptomatic and symptomatic groups were then seeded on U-shaped round-bottom 96-well cell culture plates (FisherScientific) containing cell culture media with or without stimulants. For each subject, 10^6^ PBMCs were seeded per well in triplicate for each experimental condition. Four experimental conditions consisted of PBMCs cultured in RPMI media (negative/baseline control), lipopolysaccharide from *E. coli* 0111:B4 (LPS, 100 ng/ml, Sigma-Aldrich), beta (β)-1,3-glucan derived from *Euglena* (1 µg/ml, Sigma-Aldrich), or T lymphocyte co-activators (*α*-CD3 + *α*-CD28 monoclonal antibodies, 1 µg/ml, Invitrogen). For T lymphocyte coactivation, cell culture wells were first coated with anti-CD3 antibodies for 24 hours at 4°C before PBMCs were seeded with anti-CD28 antibodies the next day. The supernatant was collected from each well after 36 hours for cytokine quantification as published (37). Cytokine quantification was performed using LEGENDplex^TM^ Human T Helper Cytokine Panels Version 2 and Human Cytokine Panel 2 (BioLegend) per manufacturer’s instructions (38). Cytokines measured in each panel included: IL-1*α*, IL-1β, IL-2, IL-4, IL-5, IL-6, IL-9, IL-10, IL-11, IL-12p40, IL-12p70, IL-13, IL-15, IL-17A, IL-17F, IL-18, IL-22, IL-23, IL-27, IL-33, IFN-*α*2, IFN-γ, GM-CSF, TNF, and TSLP.

### Statistical Analysis

Statistical analyses were performed using GraphPad Prism 8 software. For descriptive statistics, summary measurements including means (standard deviation, SD) or medians (interquartile range, IQR or percentiles) for continuous normally or non-normally distributed variables were used, respectively. The Shapiro-Wilk test with quantile-quantile plots was used to determine normal distribution. For normally distributed data, a parametric, unpaired, two-tailed Student’s t-test was used to determine significance between asymptomatic and symptomatic individuals for cytokine quantification. For non-normally distributed data, the nonparametric Mann-Whitney U test with Dunn’s multiple comparisons was used to determine significance between the two groups. For all tests performed, statistical significance was defined as \**p<0.05*, \*\**p<0.01*, \*\*\**p<0.001*, and \*\*\*\**p<0.0001*. For cytokine analysis, LEGENDplex^TM^ data analysis software suite (Qognit) was utilized to generate a standard curve for twenty-five analytes and the raw concentration values in picograms per milliliter for asymptomatic and symptomatic individuals’ PBMCs that were either stimulated or unstimulated (baseline). To account for differences in asymptomatic and symptomatic individuals’ baseline cytokine levels, raw concentrations from stimulated PBMCs were normalized to their individual raw concentrations from unstimulated conditions (RPMI media only) to calculate the linear relative fold change. Log10-base transformation of linear fold change (Log_10_[relative fold change]) was then performed before presenting medians on the heatmaps. Heatmaps were generated in R using the heatmap library. To determine associations between cytokine levels from *α*-(CD3+CD28)-stimulated PBMCs and oscillometry parameters, the degree of linear association between these two variables was assessed using the Pearson correlation coefficient (r). For LPS- and β-stimulated PBMCs’ cytokine levels not meeting normal distribution, a nonparametric Spearman correlation was performed to measure the strength of a nonlinear association (*r_rs_*) between these two variables.

## RESULTS

### Study Population Characteristics and Demographics

The cohort consisted of forty never-smoker Veterans who were previously deployed to Southwest Asia and Afghanistan and were initially recruited to the SHADE study to characterize their inhalational exposures during deployment and associations with respiratory symptoms based on the inclusion and exclusion criteria (2). They completed a questionnaire regarding deployment exposures to pollutant sources, grouped a priori into burn pit smoke, combustion particulates, vapors, gas, dusts, or fumes (VGDFs), or other toxicants categories (**Fig E1**). From the SHADE cohort, individuals were randomly selected into the LEDE study, to determine how previous exposure sources affect lung physiology, structure, and systemic inflammation biomarkers. The participants were predominantly male (n=32; 80%) and with a median age of 45 (**Table I**). At their study visit, the median body mass index (BMI) was 30.2 kg/m^2^ which was not significantly different between the groups. The total median deployment duration was 11.3 months with the study visit completed mean of 14.8 years since last deployment (**Table II**), while 25% of the cohort reported more than one deployment. The time between the SHADE visit and the follow-up visit was a median of 4 years (**Table II**). The characteristics of individuals with and without symptoms showed that the distribution of age, sex, race/ethnicity, BMI, time between study visits, years since last deployment, and deployment duration was similar between the asymptomatic and symptomatic groups.

**Table I.** Demographics of study cohort.

| **Characteristic** | **Total**<br>**n=40** | **Asymptomatic**<br>**n=24** | **Symptomatic**<br>**n=16** |
| --- | --- | --- | --- |
| **Age in Years, med (25%, 75%)** | 45 (38, 55) | 41 (38, 51) | 48 (41, 57) |
| **Sex, n (%)** |  |  |  |
| **Male** | 32 (80) | 20 (83) | 12 (75) |
| **Female** | 8 (20) | 4 (17) | 4 (25) |
| **Race and Ethnicity, n (%)** |  |  |  |
| **African American or Black** | 6 (15) | 4 (17) | 2 (13) |
| **Asian or Pacific Islander** | 2 (5) | 2 (8) | 0 (0) |
| **Multiracial or Other** | 1 (3) | 0 (0) | 1 (6) |
| **Hispanic** | 4 (10) | 1 (4) | 3 (19) |
| **White** | 31 (78) | 18 (75) | 13 (81) |
| **Body Mass Index, med (25%, 75%)** | 30.2 (26.2, 34.5) | 29.6 (26.2, 34.5) | 31.1 (25.8, 35.0) |
| All individuals are never smokers in this study. Age and percentage of individuals categorized by sex, race, and ethnicity are rounded to the nearest integer; med: median (25th percentile - 75th percentile) |  |  |  |

**Table II.**
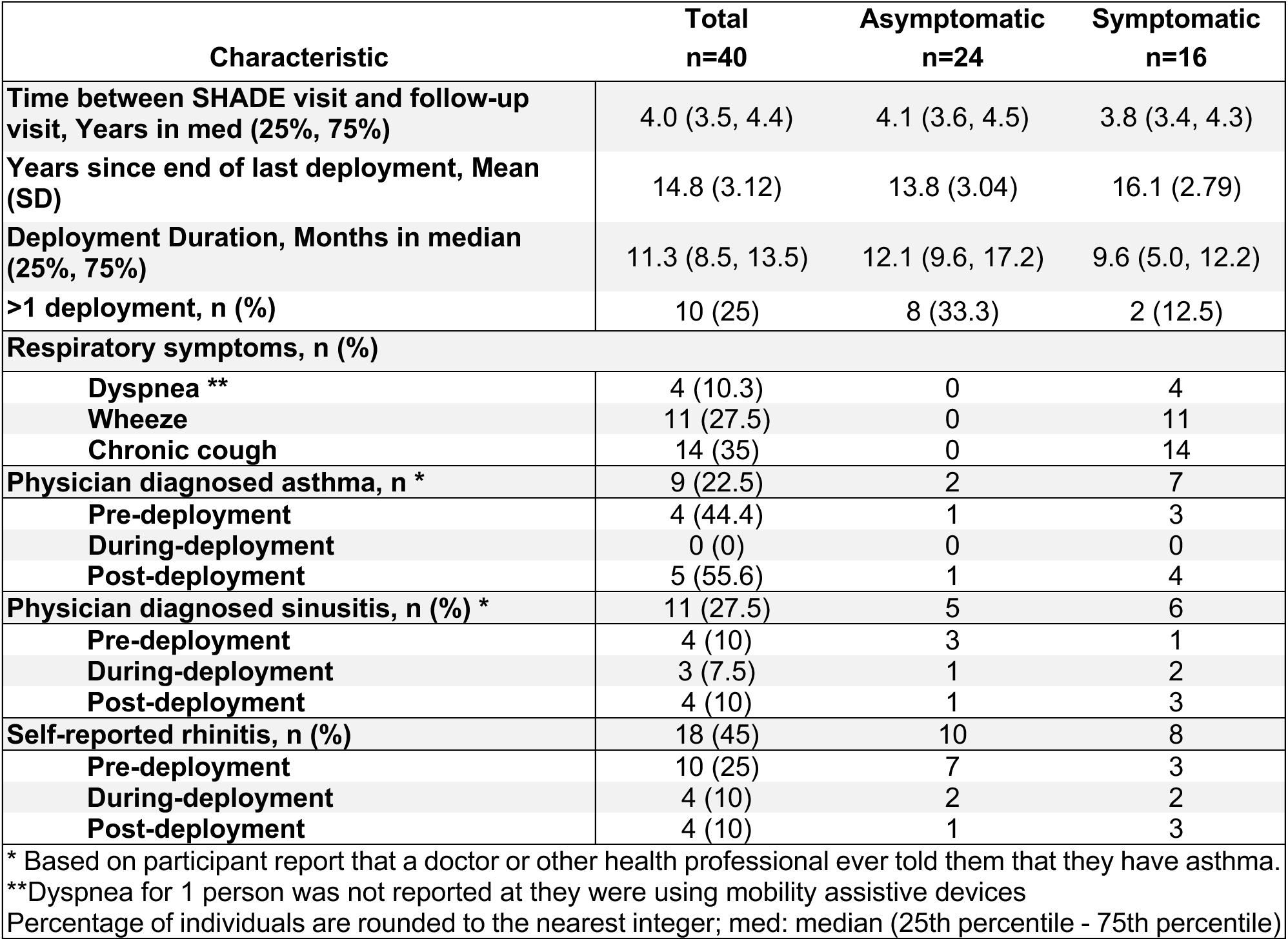
Clinical characteristics of study cohort.

| Characteristic | Total<br>n=40 | Asymptomatic<br>n=24 | Symptomatic<br>n=16 |
| --- | --- | --- | --- |
| Time between SHADE visit and follow-up visit, Years in med (25%, 75%) | 4.0 (3.5, 4.4) | 4.1 (3.6, 4.5) | 3.8 (3.4, 4.3) |
| Years since end of last deployment, Mean (SD) | 14.8 (3.12) | 13.8 (3.04) | 16.1 (2.79) |
| Deployment Duration, Months in median (25%, 75%) | 11.3 (8.5, 13.5) | 12.1 (9.6, 17.2) | 9.6 (5.0, 12.2) |
| >1 deployment, n (%) | 10 (25) | 8 (33.3) | 2 (12.5) |
| **Respiratory symptoms, n (%)** |  |  |  |
| Dyspnea ** | 4 (10.3) | 0 | 4 |
| Wheeze | 11 (27.5) | 0 | 11 |
| Chronic cough | 14 (35) | 0 | 14 |
| Physician diagnosed asthma, n * | 9 (22.5) | 2 | 7 |
| Pre-deployment | 4 (44.4) | 1 | 3 |
| During-deployment | 0 (0) | 0 | 0 |
| Post-deployment | 5 (55.6) | 1 | 4 |
| Physician diagnosed sinusitis, n (%) * | 11 (27.5) | 5 | 6 |
| Pre-deployment | 4 (10) | 3 | 1 |
| During-deployment | 3 (7.5) | 1 | 2 |
| Post-deployment | 4 (10) | 1 | 3 |
| Self-reported rhinitis, n (%) | 18 (45) | 10 | 8 |
| Pre-deployment | 10 (25) | 7 | 3 |
| During-deployment | 4 (10) | 2 | 2 |
| Post-deployment | 4 (10) | 1 | 3 |

\* Based on participant report that a doctor or other health professional ever told them that they have asthma.
\*\*Dyspnea for 1 person was not reported at they were using mobility assistive devices
Percentage of individuals are rounded to the nearest integer; med: median (25th percentile - 75th percentile)

Veterans with reported persistent respiratory symptoms (PRS) informed by dyspnea, wheezing, and chronic cough was only found in symptomatic group (**Table II**). Physician diagnosed asthma was 9 individuals in total with majority being in the symptomatic group (**Table II**). For physician diagnosed sinusitis and self-reported rhinitis, the distribution was similar between the two groups (**Table II**). There was no significant differences in the distribution of each of the three diagnosis between pre-, during- and post-deployment (**Table II**).

Most individuals (n=39; 98%) reported exposure to burn pits, combustion byproducts, other toxicants, and vapors, gas, dusts, or fumes (VGDFs) during their deployment (**Fig E1 and Table E1**). They also reported exposure to combustion sources (n=38; 95%), such as vehicle engines, generator exhaust, and those in combat or military-related, with similar percentages in both groups (ASx, n=22, 92%; Sx, n=16, 100%) (**Fig E1 and Table E1**). Both groups (ASx, n=16, 67%; Sx, n=12, 75%), reported exposure to burn pit smoke outdoors, indoors at nearby worksites or homes, or physical exertion beside a burn pit site (**Fig E1 and Table E1**). Symptomatic individuals (n=13; 81%) and asymptomatic individuals (n=14; 58%) also reported exposure to VGDF (**Fig E1 and Table E1**). Individuals’ exposure to other toxicants, such as dust storm particles, pesticides, insecticides, or repellents, was the lowest in percentage (n=21; 53%) compared to burn pits, VGDF, and combustion sources (**Fig E1 and Table E1**). Together, exposure analysis indicated that the most prevalent sources of exposure were open combustion and burn pits.

### PRS Individuals Displayed Airway Changes Consistent with Early Small Airway Dysfunction

To quantify lung function, all individuals underwent PFTs before and after bronchodilator administration to assess for changes in lung function (**Fig E2 and E3; Table E2 and E3**). There were no significant differences in percent predicted in FEV_1_, FVC, FEV_1_/FVC ratio, TLC, or DL_CO_ between the two groups pre- and post-bronchodilator (BD) administration (**Fig E2; Table E2 and E3**). Likewise, there was also no significant percent changes before and after bronchodilator administration for FEV_1_, FVC, and FEV_1_/FVC ratio (**Fig E3; Table E2 and E3**).

To further assess airway changes, oscillometry testing before and after bronchodilator administration was performed on all individuals (**Fig 1, B and C; Fig E4 and E5; Table E4 and E5**) (39). There was a significant (fourfold) increase in airway resistance detected in the symptomatic group as determined by the difference between pre-BD R_5_ and R_19_ (R_5_-R_19_) (**Fig 1, B**). The pre-BD resonant frequency (F_res_) was also significantly higher in symptomatic individuals compared to asymptomatic (**Fig 1, C**). The symptomatic group showed a trend for higher pre- and post-BD reactance area (AX), compared to the asymptomatic individuals (**Fig E4, C and G**). There was no difference observed between the groups’ respiratory system reactance (X_5_) (**Fig E4, D and H**). Additionally, CT quantification revealed no differences in percentages in low attenuation area (LAA), gas trapped, and airway wall thickness (Pi10) between the groups (**Fig E6**). Together, these findings suggest that those with PRS exhibit early alterations in the airway changes that were not detected by the PFTs or using the CT scan.

**Fig 1.**
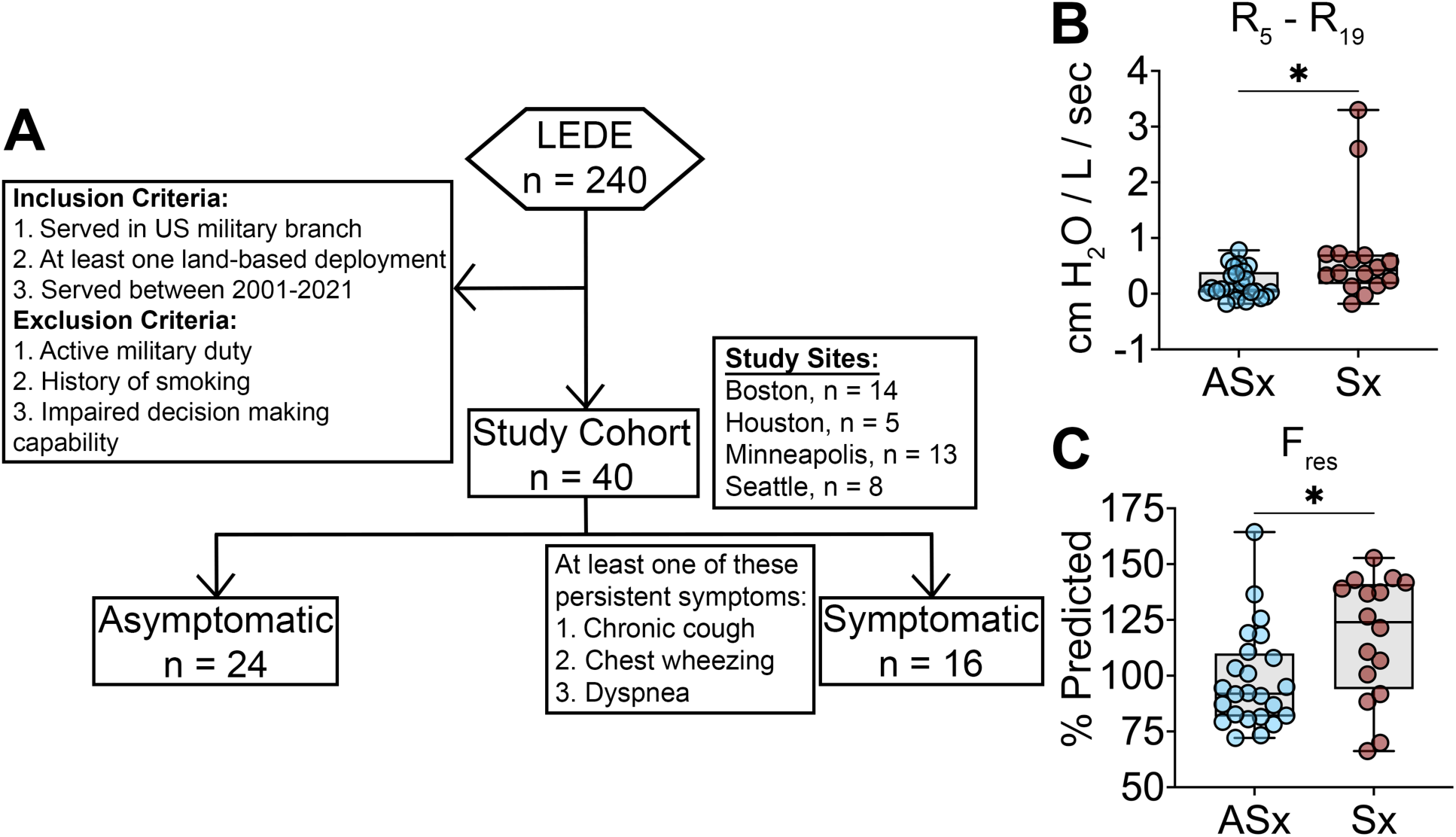
Demographics and clinical characteristics of pollutant-exposed individuals in study cohort. **(A)** Individuals with previous land-based deployment who participated in the Lung Effects of Deployment Exposure (LEDE) completed a respiratory health questionnaire. Forty non-smoker individuals with persistent (Sx, n=16) and without (ASx, n=24) respiratory symptoms were selected for analysis of oscillometry measurements before albuterol administration. Measurements included **(B)** frequency dependence of resistance (R_5_-R_19_) and **(C)** percent predicted of resonant frequency (F_res_). Medians are presented with minimum, maximum, and IQR. Significant differences between asymptomatic and symptomatic individuals were determined by Mann-Whitney U test, \**p < 0.05*.

### Differential cell counts and baseline cytokines fail to distinguish PRS individuals

We evaluated white blood cell differential cell counts for evidence of a T2 phenotype. We found no significant differences between the groups in total or relative abundance of neutrophils, lymphocytes, monocytes, eosinophils, and basophils between the groups (**Fig E7**). We next assessed differences in systemic immunity in both asymptomatic (n=24) and symptomatic (n=16) individuals by using immune cell activation of PBMCs (**Fig 2, A**). Baseline levels of a comprehensive list of cytokines that were associated with anti-inflammation, allergic/asthma response, acute inflammation, antimicrobial, and inflammasome did not show any significant differences between the groups (**Fig 2, B**). Notably, six cytokine levels (IL-11, IL-12 p40, IL-12 p70, IL-15, IL-27, and TSLP) were below the level of detection and were not included in the subsequent analyses.

**Fig 2.**
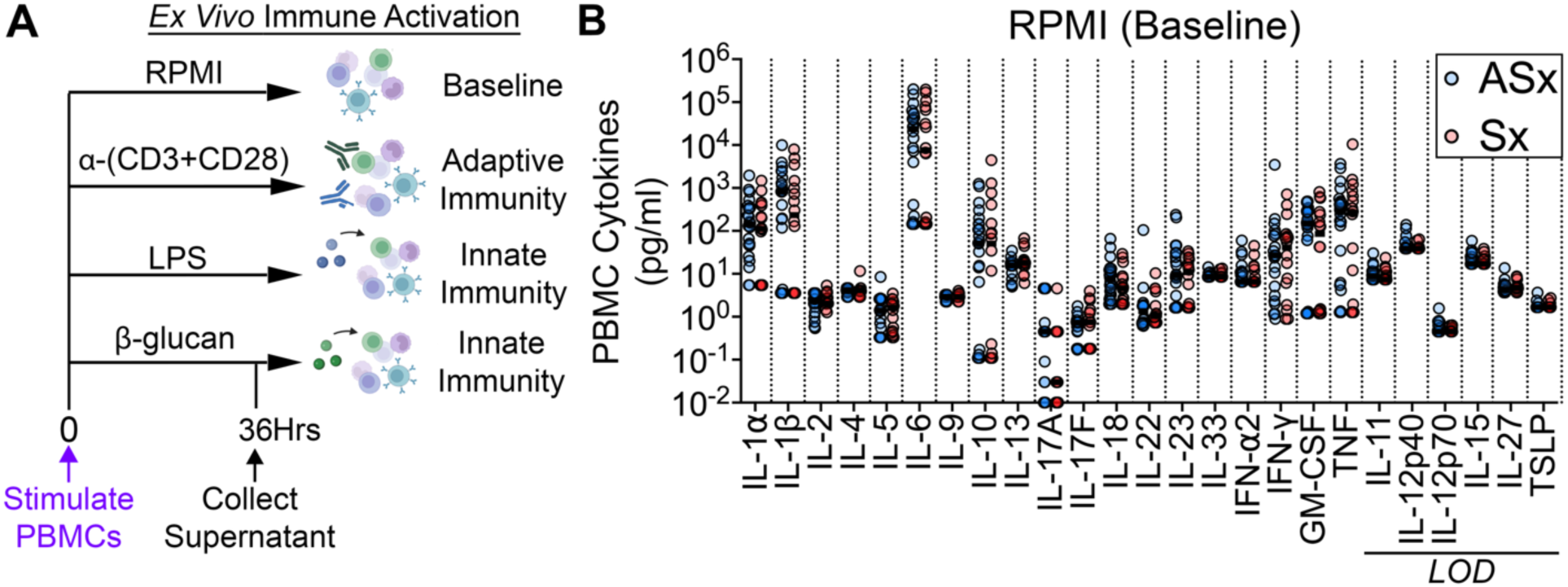
Individuals with and without persistent respiratory symptoms exhibit no pre-existing differences in cytokine profile at baseline. **(A)** Immune activation was assessed *ex vivo* using individuals’ PBMCs (10^6^ cells/well in triplicates) stimulated with vehicle (baseline control: RPMI cell culture media), T-lymphocyte coactivators (*α*-CD3 and *α*-CD28 antibodies, 1 µg/ml), lipopolysaccharide (LPS, 100 ng/ml), or beta-(β)-glucan (1 µg/ml) for 36 hours. Supernatants were then collected to measure cytokine production using the LegendPlex multiplex kit. (**B**) Individual data points of subjects from asymptomatic (ASx, n=24) and symptomatic (Sx, n=16) groups. Six cytokines did not exceed the limit of detection (LOD) as indicated. No significant differences were detected between the two groups using an unpaired Student’s t-test, \**p < 0.05*.

### Individuals with PRS exhibited increased T17 and reduced T1 immune responses

Next, we assessed altered responsiveness in systemic adaptive immunity by measuring cytokine production in response to activating T cells (**Fig 3, A; Fig E8 and E9**). We used the log10-relative fold change in each cytokine detected before and after T cell stimulating conditions. Compared to the asymptomatic group, activation of T cells in the symptomatic group exhibited a significant increase in IL-17A (**Fig 3**), while despite a trend, several other cytokines did not reach significance (**Fig E8, B**). We detected increased IL-17A, IL-17F, IL-22, and IL-23 following T cell activation in individuals with persistent symptoms (**Fig 3; Fig E8, B; and Fig E9**). Notably, we found a significant decrease in IL-6 and IFN-γ in the symptomatic group when compared to the asymptomatic (**Fig 3**). There was also a trend for increased IL-5 and IL-9 in symptomatic individuals, consistent with an altered T2 phenotype, but did not reach significance (**Fig E8, B and Fig E9**). Together, these results suggest that individuals with persistent respiratory symptoms exhibit a select immune response that is skewed towards both an enhanced proinflammatory T17 phenotype and a suppressed T1 response associated with skewed proinflammatory pathways.

**Fig 3.**
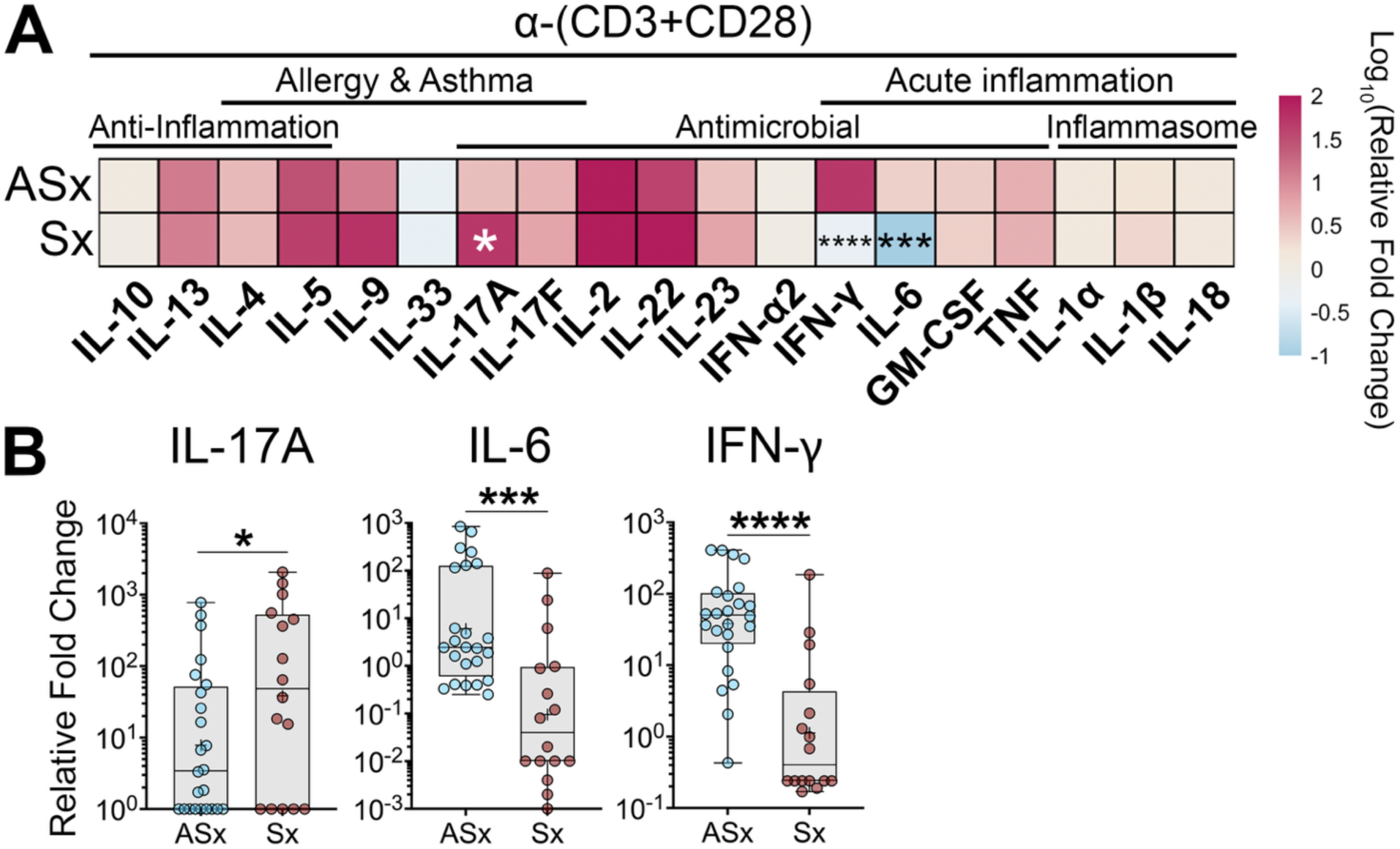
Individuals with persistent respiratory symptoms exhibited a mixed T2/T17 adaptive immune signature with reduced T1 phenotype. *Ex vivo* immune activation was assessed using individuals’ PBMCs (10^6^ cells/well) stimulated with T-lymphocyte coactivators (*α*-CD3 and *α*-CD28 antibodies, 1 µg/ml) for 36 hours before supernatant collection. Cytokine levels were measured using the LegendPlex multiplex kit. **(A)** Heatmaps represent log_10_ of medians in relative fold changes from asymptomatic (ASx, n=24) and symptomatic (Sx, n=16) individuals. Cytokines exceeding the threshold for limit of detection are displayed. **(B)** Representative panel of cytokines displayed with log_10_-transformed relative fold change. Boxplot displays median with minimum, maximum, and IQR and + represents mean. Fold change was normalized to each individual’s unstimulated PBMCs (baseline) before log_10_-transformation. Significant differences between asymptomatic and symptomatic groups determined by unpaired Student’s t-test, *\*p < 0.05, **p < 0.01, ***p < 0.001*, \*\*\*\**p < 0.0001*.

### A differential immune signature associated with altered antimicrobial response in PRS individuals

Since the PRS group displayed a skewed T17 and reduced T1 activation both associated with antimicrobial response, we next assessed the innate immune responses in the same cohort (**Fig 4; Fig E10 and Fig E11**). Upon exposure to either LPS or β-glucan, the PRS group showed a significant increase in the alarmin cytokine, IL-33 (40) (**Fig 4, A**). In the symptomatic group, there was a decrease in IL-17F and T1-associated cytokines IFN-γ and TNF specific to LPS (**Fig 4, A and Fig E10, B**). There were no significant differences in IL-1β, IL-2, IL-4, IL-6, IL-9, IL-10, IL-13, IL-17A, IL-22, IFN-*α*2, and granulocyte macrophage-colony stimulating factor (GM-CSF), while IL-23 and cytokines associated with inflammasome activation (e.g., IL-1*α* and IL-18) displayed an increased trend in the symptomatic group (**Fig E10, B**).

**Fig 4.**
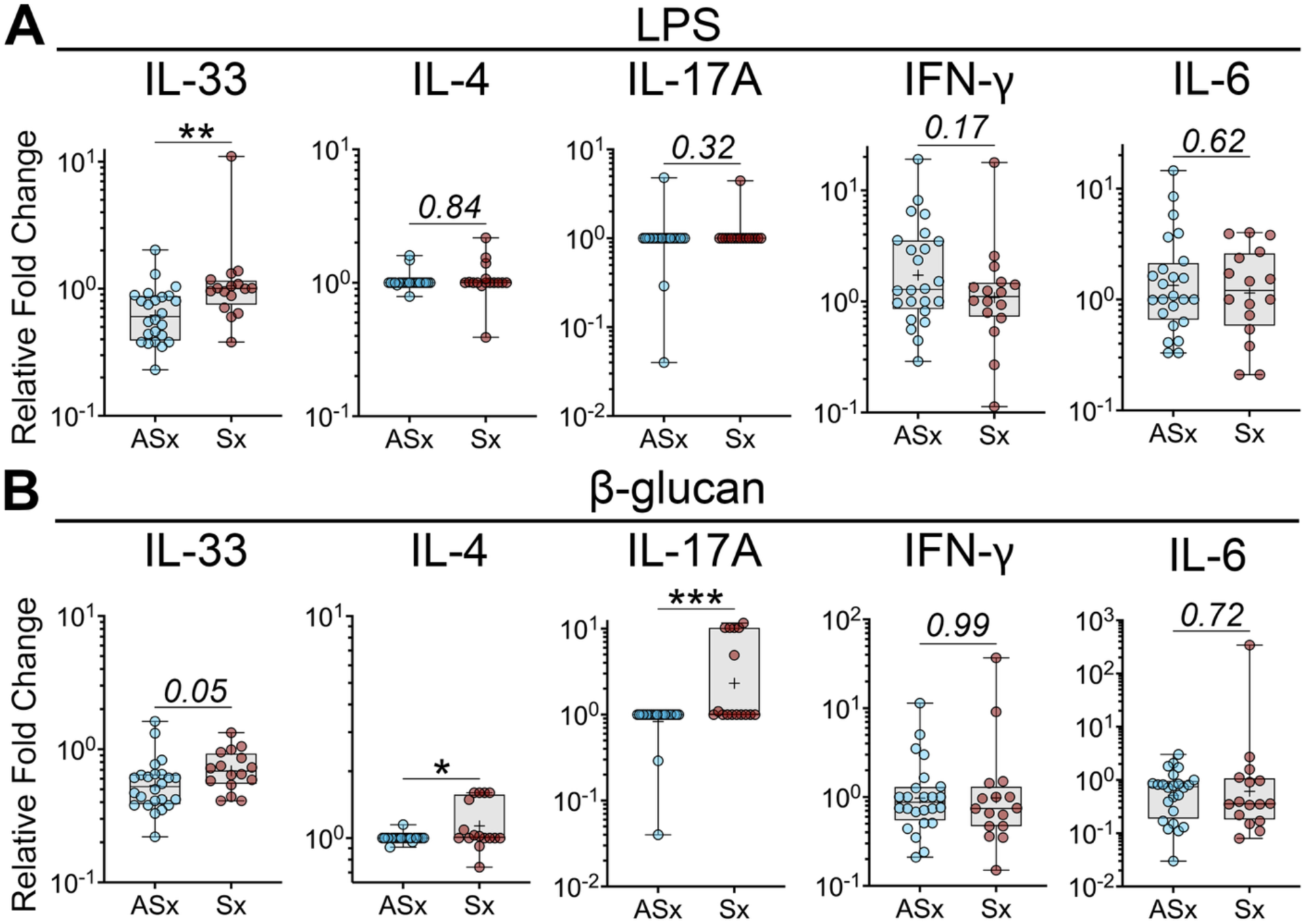
Innate immune stimulation induced a mixed T2/T17 phenotype and reduced T1-associated antimicrobial response in individuals with persistent respiratory symptoms. Immune activation was assessed by stimulating individual’s PBMCs (10^6^ cells/well) with either **(A)** LPS (100 ng/ml) or **(B)** β-glucan (1 µg/ml) for 36 hours before supernatant collection. Cytokine levels were measured using the LegendPlex multiplex kit. Representative panel of cytokines exceeding the LOD is displayed with log_10_-transformed relative fold change. Boxplot displays median with minimum, maximum, and IQR, and + represents mean. Fold change was normalized to each individual’s unstimulated PBMCs (baseline) before log_10_-transformation. Significant differences between asymptomatic (ASx, n=24) and symptomatic (Sx, n=16) groups determined by unpaired Student’s t-test, *\*p < 0.05, **p < 0.01, ***p < 0.001*, \*\*\*\**p < 0.0001*.

Notably, β-glucan stimulation caused a significant increase in both IL-4 and IL-17A in the PRS group (**Fig 4, B**). While stimulation with β-glucan caused no differences in many cytokines (**Fig E11, B**), we detected a trend for decreased inflammatory myeloid-associated cytokines GM-CSF, IL-6, and TNF in the PRS group (**Fig 4, B and Fig E11, B**). In addition, inflammasome-associated cytokines IL-1*α*, IL-1β, and IL-18 also showed a decreasing trend in the PRS group (**Fig E11, B**). Together, exposure to LPS or β-glucan revealed a differential immune signature associated with altered antimicrobial response in PRS group.

### Mixed T2/T17 cytokine activation correlates with oscillometry measurements of early small airway dysfunction

Given our earlier findings that showed significant differences in oscillometry measurements between the two groups, we next determined if there is a relationship between cytokine levels following T cell or innate immune activation and peripheral airway function (**Fig 5; Fig E12; and Table III**).

**Fig 5.**
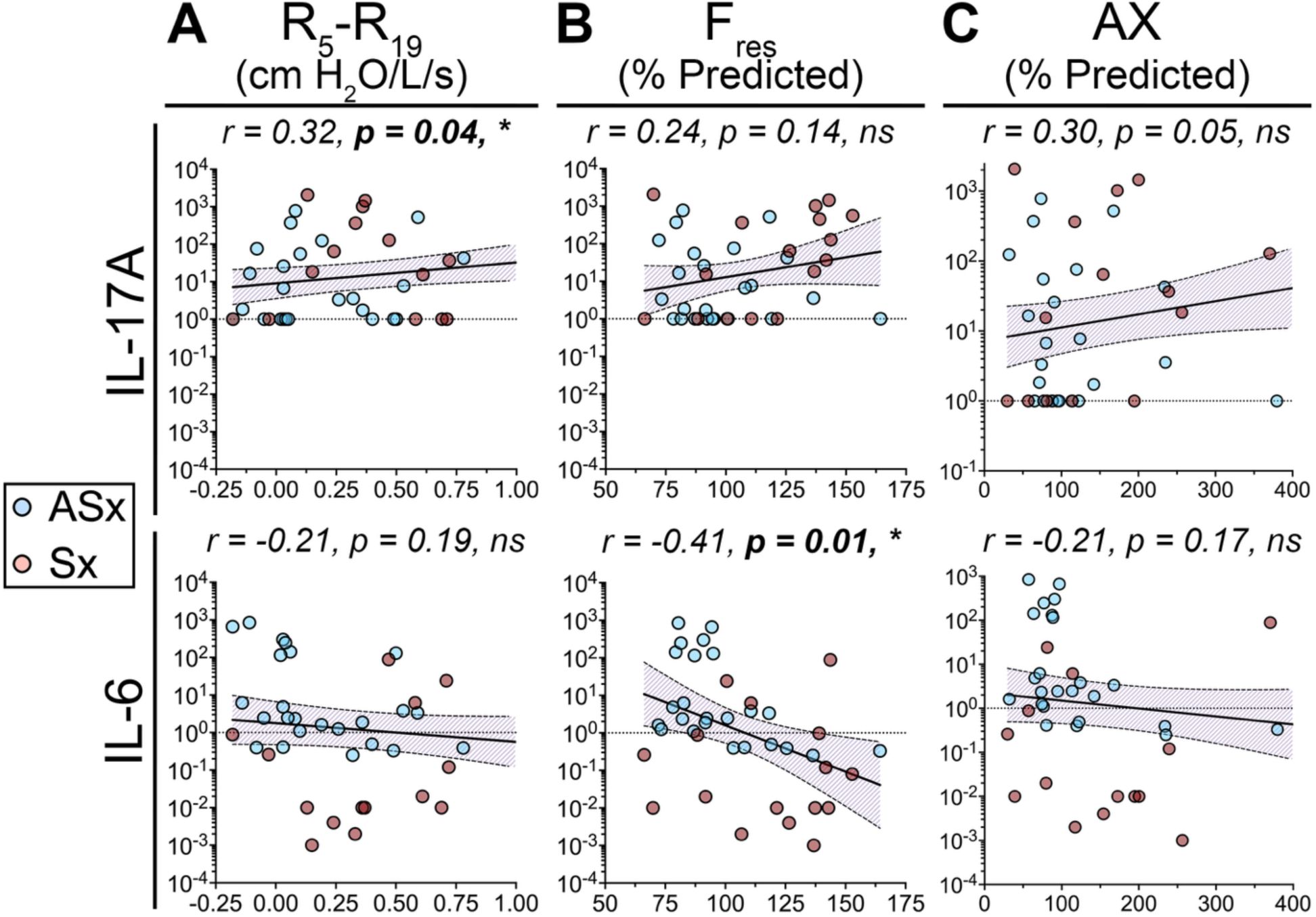
T17-associated cytokines positively correlated with airway abnormalities, consistent with early small airway disease. Correlation analysis was performed in pollutant-exposed individuals to determine the correlation strength (*r*) between the log_10_-fold change of IL-17A and IL-6 in *α*-(CD3+CD28)-stimulated PBMCs with (**A**) small airway resistance (R_5_-R_19_), (**B**) resonant frequency (F_res_), and (**C**) area of reactance (AX). Each data point represents each asymptomatic (n = 24, blue dots) or symptomatic (n = 16, red dots) individual. The dashed line represents a linear fold change of 1, which represents unstimulated PBMCs. The shaded region represents error bands within 95% confidence interval. Pearson correlation analysis was performed to determine significance. *\*p < 0.05, **p < 0.01*.

**Table III.** Bivariate analysis assessing innate immune stimulation and airway changes in pollutant-exposed individuals.

|  | **R<sub>5</sub>-R<sub>19</sub>**<br>**(cm H<sub>2</sub>O/L/sec)** |  | **F<sub>res</sub>**<br>**(% Predicted)** |  | **AX**<br>**(% Predicted)** |  |
| --- | --- | --- | --- | --- | --- | --- |
|  | *r<sub>s</sub>* | *p-value* ** | *r<sub>s</sub>* | *p-value* ** | *r<sub>s</sub>* | *p-value* ** |
| **LPS stimulation\*** |  |  |  |  |  |  |
| **IL-17A** | -0.07 | 0.66 | -0.07 | 0.64 | -0.08 | 0.61 |
| **IL-4** | 0.10 | 0.52 | 0.04 | 0.81 | 0.14 | 0.40 |
| **IL-33** | -0.01 | 0.97 | 0.02 | 0.88 | -0.05 | 0.74 |
| **β-glucan stimulation\*** |  |  |  |  |  |  |
| **IL-17A** | 0.46 | **0.003** | 0.45 | **0.003** | 0.40 | **0.01** |
| **IL-4** | 0.47 | **0.002** | 0.24 | 0.12 | 0.23 | 0.15 |
| **IL-33** | 0.27 | 0.09 | 0.31 | 0.05 | 0.34 | **0.03** |

\*Cytokine levels were measured in log<sub>10</sub>(Relative Fold Change), where fold change post-stimulation with either LPS (100 ng/ml) or β-glucan (1 μg/ml) was normalized to unstimulated PBMCs (baseline) before log<sub>10</sub>-transformation.
\*\*Spearman correlation was performed to determine correlation strength (*r<sub>s</sub>*) between log<sub>10</sub>(Relative Fold Change) of IL-17A, IL-4, and IL-33 produced by stimulated PBMCs with frequency dependence of resistance (R<sub>5</sub>-R<sub>19</sub>), resonant frequency (F<sub>res</sub>), and area of reactance (AX). \**p*<0.05, \*\**p*<0.01

We found a significant positive correlation between T cell activation-induced IL-5 (r = 0.33, *p = 0.03*), IL-9 (r = 0.33, *p = 0.03*), and IL-17A (r = 0.32, *p = 0.04*) with R_5_-R_19_ (**Fig 5, A and Fig E12, A**). Although not significant, there was a trend for negative correlation between IL-6 and R_5_-R_19_ (r = -0.21; *p = 0.20*) (**Fig 5, A**). Similarly, we found a positive correlation between IL-9 (r = 0.30, *p = 0.05*) induction and a significant negative correlation for IL-6 (r = -0.41, *p = 0.01*) with resonant airway frequency (**Fig 5, B and Fig E12, B**). Furthermore, IL-5 (r = 0.31, *p = 0.04*), IL-9 (r = 0.32, *p = 0.04*), and IL-17A (r = 0.30, *p = 0.05)* production positively correlated with reactance area (AX), while IL-6 (r = - 0.22, *p = 0.18)* had a negative correlation (**Fig 5, C and Fig E12, C**). These findings are consistent with a T cell-mediated induction of T2-associated IL-5 and IL-9 and T17- associated IL-17A response correlating with increased small airway resistance, airway stiffness, and decreased small airway compliance. Notably, a decrease in IL-6 (T1) negatively correlated with the same specific lung mechanics. Since IL-5, IL-9, and IL-17A from T-cell activation showed significant correlations across the three oscillometry measurements, we next assessed if there are associations between immune activation by pathogen-associated molecular patterns, LPS, and β-glucan and small airways alterations (**Table III**). No significant associations were detected between the selected cytokines and the three oscillometry measurements for LPS-stimulated immune cells (**Table III**). However, significant positive non-linear associations were detected between T2-associated IL-4 with R_5_-R_19_ (*r_s_* = 0.47, *p = 0.002*) for β-glucan-stimulated immune cells (**Table III**). IL-17A showed the same significant positive associations with R_5_-R_19_ (r_s_ = 0.46, *p = 0.003*), F_res_ (r_s_ = 0.45, *p = 0.003)*, and AX (r_s_ = 0.40, *p = 0.01*) (**Table III**). IL-33 displayed positive associations with F_res_ (r_s_ = 0.31, *p = 0.05*) and AX (r_s_ = 0.34, *p = 0.03*) (**Table III**). Together, these analyses suggest the potential role of IL-5, IL-9, and IL-17A cytokines associated with T2 and T17 pathways in small airway dysfunction that indicates an early activation of the pathogenic pathway in exposed individuals with persistent symptoms. Furthermore, IL-17A, IL-4, and IL-33, associated with the same T2/T17 pathways, may also contribute to small airway dysfunction and altered antimicrobial response that is pathogen-specific.

## DISCUSSION

This study identifies a potential early respiratory phenotype in pollutant-exposed individuals with PRS, characterized by oscillometric evidence of small-airway dysfunction and stimulus-dependent systemic immune dysregulation. The correlation between these abnormalities suggests a biological link that is not captured by conventional spirometry, chest CT, resting cytokine levels, or leukocyte counts. Although occupational and environmental air pollutants are well-established contributors to respiratory disease (41–43), predicting which exposed individuals will develop persistent symptoms remains challenging. Sensitive physiological and immune assessments may therefore help detect early disease processes before abnormalities emerge on standard pulmonary function tests. We examined deployed Veterans with documented inhalational exposures, with or without persistent respiratory symptoms. Despite the modest cohort size, stimulus-induced mixed cytokine responses correlated with oscillometric evidence of small-airway dysfunction across pollutant-exposed participants. These findings support an association between altered systemic immune responsiveness and early small-airway dysfunction in this population.

Immune activation is tightly regulated and must adapt to diverse environmental stimuli, including pollutants, allergens, and microbes (12, 44). This response requires coordinated activity among T and B lymphocytes, myeloid cells, and innate lymphoid cells, which collectively shape T1, T2, and T17 immunity (12). Effective host defense depends on sufficient cytokine production to eliminate pathogens while limiting excessive inflammation and tissue injury. Cytokines integrate effector and regulatory signals that initiate, sustain, and resolve systemic immune responses; disruption of this balance can contribute to chronic pulmonary and systemic disease (44).

In PBMCs from Veterans with documented inhalational exposures, baseline cytokine levels did not differ between the groups, arguing against constitutive systemic cytokine elevation in PRS. After adaptive immune activation, however, the PRS group exhibited a distinct mixed T2/T17 signature accompanied by reduced T1 responses. Although our *ex vivo* experiments measured cytokine production rather than cellular sources, the increased IL-17A, IL-17F, IL-1β, IL-22, and IL-23 suggests an inducible T17-centered inflammatory axis (12). The study opens this line of investigation in future studies to identify the responsible cell populations in PBMC among the diverse immune subsets, including T and B lymphocytes, myeloid cells, and innate lymphoid cells (45, 46).

Harmful inhaled stimuli can promote Th17-cell differentiation and IL-17A production (47). In asthma, T17-polarized airway inflammation is associated with severe, steroid-resistant disease in both mice and humans (48, 49). Consistent with activation of this pathway, T-cell stimulation in participants with PRS increased IL-17A, IL-17F, IL-22, and IL-23, cytokines implicated in airway, asthmatic, and allergic inflammation (50). The concurrent reduction in T1 responses that we found in this study suggests a shift in systemic adaptive immunity toward T17 dominance. IL-6 can promote early Th17 differentiation through classical signaling via membrane-bound IL-6Rα and gp130 or through trans-signaling mediated by soluble IL-6Rα complexes in the tissue microenvironment (51, 52). Dysregulation of these signaling pathways may contribute to systemic inflammatory disease (53).

IL-17A correlated positively, whereas IL-6 correlated negatively, with oscillometric abnormalities, suggesting potential uncoupling of the IL-6/IL-17 axis in pollutant-exposed individuals with PRS. IL-6 signaling is normally constrained by SOCS1 and SOCS3, which inhibit JAK pathways and gp130, respectively, through negative-feedback mechanisms (54). In contrast, IL-22 and IL-23 can sustain Th17 responses independently of IL-6 (55). Thus, increased IL-22 and IL-23 production in the setting of reduced IL-6 and IFN-γ may reflect altered intracellular signaling and immune-cell crosstalk among T cells, myeloid cells, and innate lymphoid cells in PRS. Future *ex vivo* and *in vivo* studies integrating transcriptomic and epigenetic analyses should determine which IL-6 signaling pathways are suppressed in specific immune-cell populations after airborne pollutant exposure.

Persistent exposure to airborne pollutants has been associated with increased susceptibility to respiratory infection (56). Because participants with PRS exhibited reduced T1-associated antimicrobial responses after T-cell activation, we examined systemic immune responses to LPS and β-glucan, which model recognition of Gram-negative bacterial and fungal stimuli, respectively. Both stimuli increased production of IL-33, a potent alarmin, in the symptomatic group. During allergic inflammation and helminth infection, IL-33 released from injured alveolar epithelial cells activates ST2-expressing ILC2s and mast cells, promoting Th2-cell production of IL-4, IL-5, and IL-13 and recruitment of eosinophils, basophils, and alternatively activated macrophages (57). Airborne pollutants, including diesel exhaust particles that contribute to PM_2.5_, can also induce IL-33 and promote pathogenic mixed T2/T17 responses that enhance allergic airway hyperresponsiveness *in vivo* (58). Although we did not directly expose PBMCs to particulate matter, participants with PRS exhibited a mixed T2/T17 phenotype, with IL-33 induction particularly evident after β-glucan stimulation. Together, these findings suggest that pollutant-associated immune reprogramming may alter antimicrobial responses and increase susceptibility to respiratory infection in at-risk populations (56, 59).

The small airways are considered the lung’s “quiet zone,” where obstructive diseases such as COPD and asthma may develop before conventional pulmonary function tests become abnormal (60). Oscillometry is increasingly used alongside spirometry to detect small-airway obstruction (61, 62). In our symptomatic group, higher R_5_-R_19_, F_res_, and area of reactance indicated increased small-airway resistance and stiffness with reduced compliance. The absence of group differences on standard pulmonary function testing and quantitative CT may reflect the cohort’s relatively young age and never-smoking status. Larger longitudinal studies are needed to determine whether oscillometry can predict progression to defined obstructive lung disease. Complementary studies should also investigate the transcriptional and epigenetic effects of direct pollutant exposure to identify mechanisms that drive chronic pulmonary and systemic disease.

## AUTHOR CONTRIBUTIONS

Supervision: F.K. Conceptualization: A.M.M. and F.K. Data curation: M.S. and D.J. Formal analysis: A.M.M., C.H.W, E.G., V.S.F., D.T.W., R.S.J.E., and F.K. Funding acquisition: C.H.W, D.B.C., F.K., and A.M.M. Investigation: A.M.M, M.S., C.H.W., E.G., V.S.F, R.S.J.E., L.S., J.L., S.P.M., I.M.M., D.J., D.T.W., and F.K. Methodology: A.M.M, M.S., C.H.W., E.G., V.S.F, D.J., R.S.J.E., D.T.W., and F.K. Project administration: M.S., C.H.W., D.J., L.S., and F.K. Resources: M.S., C.H.W., E.G., V.S.F, D.J., R.S.J.E., D.T.W., L.S. D.B.C., and F.K. Software: M.S., R.S.J.E., and D.J.. Validation: M.S. and D.J. Visualization: A.M.M. and F.K. Writing - original draft: A.M.M and F.K. Writing - review & editing: A.M.M, M.S., C.H.W., E.G., V.S.F, D.B.C., D.J., D.T.W., R.S.J.E., and F.K.

## ACKNOWLEDGEMENTS

This work was supported by the Department of Veterans Affairs (VA) Office of Research & Development (ORD) Cooperative Studies Program Study #595 BLR&D I01 BX004619, I01 BX004633, 01 BX004609-01A1 from PACT Act Toxic Exposure Funds through the VA ORD Military Exposures Research Program. Additional funding support by NIH 1R01 HL177930, and training grant 1T32HL170991-01 and the Burroughs Wellcome Fund Postdoctoral Enrichment Program 1 - 1617697. Microsoft Word Copilot was used to assist with clarity and grammar.

## DATA AVAILABILITY STATEMENT

The datasets generated and/or analyzed during the current study are not publicly available but are available for the corresponding author on reasonable request through established VA data access mechanisms (e.g., via a VA-approved Data Use Agreement and review by the relevant VA oversight bodies). Qualified researchers may obtain access subject to VA regulatory approval, ensuring protection of Veteran privacy while enabling scientific verification.

## ABBREVIATIONS

ACK: Ammonium-chloride-potassium
ASx: Asymptomatic
ATS: American Thoracic Society
AX: Area of reactance
β-glucan: Beta-glucan
BD: Bronchodilator
BMI: Body mass index
CARAS: Combined allergic rhinitis and asthma syndrome
COPD: Chronic Obstructive Pulmonary Disease
CT: Computed tomography
DL_CO_: Diffusing capacity for carbon monoxide
DMSO: Dimethyl sulfoxide
EDTA: Ethylenediaminetetraacetic acid
ERS: European Respiratory Society
FBS: Fetal bovine serum
FEV_1_: Forced expiratory volume in 1 second
F_res_: Resonant frequency
FVC: Forced vital capacity
IFN-γ: Interferon-gamma
IL: Interleukin
ILCs: Innate lymphoid cells
ILDs: Interstitial lung diseases
IQR: Interquartile range
JAK: Janus-activated kinase
LAA: Low attenuation area
LEDE: Lung Effects of Deployment Exposure
LOD: Limit of detection
LPS: Lipopolysaccharide
M: Molar (mol/L)
nCB: Nanocarbon black
PAH: Polyaromatic hydrocarbons
PBMCS: Peripheral blood mononuclear cells
PBS: Phosphate-buffered saline
PFTs: Pulmonary function tests
Pi10: Internal perimeter of 10 mm (airway thickness)
PM_2.5_: Particulate matter under 2.5 microns in diameter
PRS: Persistent respiratory symptoms
r: Correlation coefficient
r_s_: Strength of a non-linear association
R_5_: Total airway resistance at 5 Hz
R_19_: Central airway resistance at 19 Hz
R_5_-R_19_: Frequency dependence of resistance (small [peripheral] airway resistance)
RPMI: Roswell Park Memorial Institute
SD: Standard deviation
SHADE: Service and Health Among Deployed Veterans
SOCS: Suppressor of Cytokine Signaling
Sx: Symptomatic
STAT3: Signal Transducer and Activator of Transcription 3
Th: T helper cells
T1: Type 1 or Th1 response
T2: Type 2 or Th2 response
T17: Type 17 or Th17 response
TLC: Total lung capacity
VA: Veteran Affairs
VGDFs: Vapors, gases, dusts, and fumes
WBC: White blood cell
X_5_: Reactance at 5 Hz

## Notes

CONFLICTS OF INTEREST: All authors declare no relevant financial disclosure of interests.

### Competing Interest Statement

The authors have declared no competing interest.

