## Supplemental Materials for "A Mixed T2/T17-Associated Systemic Immune Signature Links Airborne Pollutant Exposure to Persistent Respiratory Symptoms"

### List of Supplementary Tables

**Supplementary Table E1.** Number and percent of individuals exposed to deployment-related airborne pollution sources

**Supplementary Table E2.** Distribution of pulmonary function values and percent with abnormal results in all individuals

**Supplementary Table E3.** Distribution of pulmonary function values in asymptomatic and persistent respiratory symptoms individuals

**Supplementary Table E4.** Distribution of oscillometry measurements in all individuals

**Supplementary Table E5.** Percent predicted and percent change of oscillometry measurements in asymptomatic and persistent respiratory symptoms individuals

### List of Supplementary Figures

**Supplementary Figure E1.** Percentage of individuals exposed to deployment-related airborne pollutant sources.

**Supplementary Figure E2.** Pulmonary function tests assessed in pollutant-exposed asymptomatic and symptomatic individuals before and after bronchodilator (BD) administration.

**Supplemental Figure E3.** Percent change of each pulmonary function tests between pre- and post-albuterol administration in pollutant-exposed asymptomatic and symptomatic individuals.

**Supplemental Figure E4.** Distribution of oscillometry measurements in asymptomatic and symptomatic individuals.

**Supplemental Figure E5.** Percent change of each oscillometry measurement between pre- and post-albuterol administration in pollutant-exposed asymptomatic and symptomatic individuals.

**Supplemental Figure E6.** Computed tomography and airway thickness assessment exhibited no differences in symptomatic compared to asymptomatic group.

**Supplemental Figure E7.** Neutrophils and lymphocytes comprise majority of white blood cells in pollutant-exposed individuals.

**Supplemental Figure E8.** Adaptive immune activation of PBMCs from pollutant-exposed individuals.

**Supplemental Figure E9.** Symptomatic individuals exhibited a mixed T2 and T17 with reduced T1 phenotype.

**Supplemental Figure E10.** PBMCs from pollutant-exposed individuals were stimulated with LPS.

**Supplemental Figure E11.** PBMCs from pollutant-exposed individuals were stimulated with β-glucan.

**Supplemental Figure E12.** T2-associated cytokines positively correlated with increased small airway resistance and decreased compliance in pollutant-exposed individuals.

**Supplementary Table E1.** Number and percent of individuals exposed to deployment-related airborne pollution sources

| Number with exposure, n (%) | Total n=40 | Asymptomatic n=24 | Symptomatic n=16 |
|---|---|---|---|
| **Any exposure** | 39 (98) | 23 (96) | 16 (100) |
| **Burn pits** | 28 (70) | 16 (67) | 12 (75) |
| **Combustion particulates** | 38 (95) | 22 (92) | 16 (100) |
| **Military-related VGDF *** | 27 (68) | 14 (58) | 13 (81) |
| **Other toxicants** | 21 (53) | 10 (42) | 11 (69) |

All individuals are never smokers in this study. Percentage of individuals are rounded to the nearest integer.
* VGDF: Vapors, gases, dusts, or fumes

**Supplementary Table E2.** Distribution of pulmonary function values and percent with abnormal results in all individuals

| Pulmonary Function Measure | Mean (SD) | Median (25%, 75%) | Abnormal* n | Min, Max |
|---|---|---|---|---|
| **Pre-albuterol % predicted **** | | | | |
| **$FEV_1$** | 103.7 (14.12) | 104.7 (95.8, 114.9) | 2 | 69.0, 124.9 |
| **FVC** | 107.0 (11.56) | 107.7 (100.2, 115.8) | 1 | 74.5, 126.7 |
| **$FEV_1$ / FVC** | 96.9 (7.50) | 98.0 (93.4, 101.5) | 1 | 64.0, 109.5 |
| **Post-albuterol % predicted** | | | | |
| **$FEV_1$** | 107 (13.4) | 110 (97.5, 118.0) | - | 77.0, 129.0 |
| **FVC** | 106 (11.1) | 107 (99.8, 116) | - | 81.4, 128.0 |
| **$FEV_1$ / FVC** | 100.7 (7.2) | 101.8 (98.2, 104.7) | - | 69.1, 111.5 |
| **TLC ***** | 94.3 (9.5) | 95.5 (86.9, 101.0) | - | 74.2, 115.0 |
| **% change ****** | | | | |
| **$FEV_1$** | 3.4 (3.70) | 2.7 (1.2, 5.7) | - | -2.4, 12.1 |
| **FVC** | -0.5 (3.19) | -0.6 (-2.4, 0.9) | - | -6.2, 12.9 |
| **$FEV_1$ / FVC** | 3.8 (2.3) | 3.2 (2.1, 5.2) | - | -0.2, 9.0 |
| **Pre-albuterol raw values **** | | | | |
| **$FEV_1$, L** | 3.7 (0.8) | 3.7 (3.3, 4.3) | - | 1.9, 5.8 |
| **FVC, L** | 4.8 (1.0) | 4.8 (4.2, 5.4) | - | 2.5, 7.1 |
| **$FEV_1$ / FVC, ratio** | 0.8 (0.1) | 0.8 (0.8, 0.8) | - | 0.5, 0.9 |
| **Post-albuterol raw values** | | | | |
| **$FEV_1$, L** | 3.9 (0.8) | 3.8 (3.3, 4.4) | - | 2.3, 5.7 |
| **FVC, L** | 4.8 (0.9) | 4.8 (4.2, 5.4) | - | 2.9, 6.9 |
| **$FEV_1$ / FVC, ratio** | 0.8 (0.1) | 0.8 (0.8, 0.8) | - | 0.5, 0.9 |
| **TLC ***, L** | 6.5 (1.2) | 6.5 (5.6, 7.4) | - | 4.0, 9.3 |
| **$DL_{CO}$ ***, mL/min/mmHg** | 27.8 (5.9) | 27.6 (23.8, 31.5) | - | 16.4, 43.6 |

* Abnormal clinical results identified was based on number of values that are at or below 5th percentile for % predicted - $FEV_1$, FVC, and $FEV_1$/FVC.

** Percent predicted and raw values for spirometry measurements except total lung capacity and diffusing capacity for carbon monoxide values were taken before subject was given albuterol (pre-bronchodilator).

*** Total lung capacity and diffusing capacity for carbon monoxide values are only taken post-bronchodilator.

**** Percent change in value for $FEV_1$, FVC, and $FEV_1$/FVC was calculated by taking the difference between precent predicted post-bronchodilator and pre-bronchodilator.

Abbreviations: $FEV_1$: Forced expiratory volume in 1 second; FVC: Forced vital capacity; TLC, Total lung capacity; $DL_{CO}$: Diffusing capacity for carbon monoxide

**Supplementary Table E3.** Distribution of pulmonary function values in asymptomatic and persistent respiratory symptoms individuals

| | Total n=40 | Asymptomatic n=24 | Symptomatic n=16 | |
|---|---|---|---|---|
| **Pulmonary Function Measure** | Mean (SD)<br>Median (25%, 75%) | Mean (SD)<br>Median (25%, 75%) | Mean (SD)<br>Median (25%, 75%) | *p-value* * |
| **Pre-albuterol % predicted** ** | | | | |
| **$FEV_1$** | 103.7 (14.12)<br>104.7 (95.8, 114.9) | 105.0 (13.61)<br>107.3 (94.9, 115.8) | 101.7 (15.08)<br>103.4 (99.8, 112.4) | *0.47* |
| **FVC** | 107.0 (11.56)<br>107.7 (100.2, 115.8) | 106.9 (11.29)<br>107.4 (98.1, 115.8) | 107.0 (12.33)<br>107.7 (103.5, 117.0) | *0.97* |
| **$FEV_1$ / FVC** | 96.9 (7.50)<br>98.0 (93.4, 101.5) | 98.1 (5.90)<br>99.5 (94.8, 102.0) | 95.1 (9.35)<br>95.6 (93.0, 101.0) | *0.23* |
| **Post-albuterol % predicted** | | | | |
| **$FEV_1$** | 107.0 (13.4)<br>110 (97.5, 118.0) | 108.0 (13.5)<br>110.0 (102.0, 116.0) | 106.0 (13.5)<br>110.0 (102.0, 116.0) | *0.67* |
| **FVC** | 106.0 (11.1)<br>107.0 (99.8, 116) | 106.0 (11.6)<br>106.0 (98.9, 116.0) | 107.0 (10.7)<br>109.0 (103.0, 114.0) | *0.83* |
| **$FEV_1$ / FVC** | 100.7 (7.2)<br>101.8 (98.2, 104.7) | 102.0 (5.3)<br>102.0 (99.4, 105.0) | 99.4 (9.5)<br>110.0 (96.7, 105.0) | *0.37* |
| **TLC** *** | 94.3 (9.5)<br>95.5 (86.9, 101) | 94.0 (8.9)<br>95.9 (87.2, 101.0) | 94.6 (10.6)<br>95.5 (86.2, 102.0) | *0.85* |
| **% Change** **** | | | | |
| **$FEV_1$** | 3.4 (3.70)<br>2.7 (1.2, 5.7) | 2.8 (3.34)<br>2.1 (1.0, 5.0) | 4.3 (4.14)<br>3.1 (1.4, 7.5) | *0.24* |
| **FVC** | -0.5 (3.19)<br>-0.6 (-2.4, 1.0) | -0.8 (2.30)<br>-0.4 (-2.3, 1.0) | -0.2 (4.25)<br>-1.4 (-2.5, 1.5) | *0.54* |
| **$FEV_1$ / FVC** | 3.8 (2.3)<br>3.2 (2.1, 5.2) | 3.4 (2.1)<br>2.8 (2.1, 4.6) | 4.2 (2.6)<br>4.6 (1.8, 5.5) | *0.29* |
| **Pre-albuterol raw values** ** | | | | |
| **$FEV_1$, L** | 3.7 (0.8)<br>3.7 (3.3, 4.3) | 3.9 (0.8)<br>4.0 (3.3, 4.4) | 3.5 (0.8)<br>3.6 (3.0, 3.9) | *0.07* |
| **FVC, L** | 4.8 (1.0)<br>4.8 (4.2, 5.4) | 5.0 (1.0)<br>5.1 (4.2, 5.5) | 4.5 (1.0)<br>4.6 (3.9, 4.9) | *0.19* |
| **$FEV_1$ / FVC, ratio** | 0.8 (0.1)<br>0.8 (0.8, 0.8) | 0.8 (0.0)<br>0.8 (0.8, 0.8) | 0.8 (0.1)<br>0.8 (0.8, 0.8) | *0.18* |
| **Post-albuterol raw values** | | | | |
| **$FEV_1$, L** | 3.9 (0.8)<br>3.8 (3.3, 4.4) | 4.0 (0.8)<br>4.1 (3.5, 4.6) | 3.6 (0.7)<br>3.7 (3.0, 4.1) | *0.09* |
| **FVC, L** | 4.8 (0.9)<br>4.8 (4.2, 5.4) | 4.9 (0.9)<br>5.0 (4.2, 5.5) | 4.5 (0.9)<br>4.5 (3.9, 4.9) | *0.20* |
| **$FEV_1$ / FVC, ratio** | 0.8 (0.1)<br>0.8 (0.8, 0.8) | 0.8 (0.0)<br>0.8 (0.8, 0.8) | 0.8 (0.1)<br>0.8 (0.8, 0.8) | *0.26* |
| **TLC ***, L** | 6.5 (1.2)<br>6.2 (5.6, 7.4) | 6.6 (1.0)<br>6.3 (5.7, 7.4) | 6.3 (1.4)<br>6.0 (5.4, 7.3) | *0.51* |
| **$DL_{CO}$ ***, mL/min/mmHg** | 27.8 (5.9)<br>27.6 (23.8, 31.5) | 28.8 (5.6)<br>29.0 (24.4, 31.8) | 26.4 (6.1)<br>25.7 (21.7, 29.3) | *0.20* |

* Significant differences between asymptomatic and symptomatic groups determined by unpaired Student t-test ($p < 0.05$).
** Percent predicted and raw values for spirometry measurements except total lung capacity and diffusing capacity for carbon monoxide value were taken before subject was given albuterol (pre-bronchodilator).
*** Total lung capacity and diffusing capacity for carbon monoxide value is post-bronchodilator.
**** Percent change in value for $FEV_1$, FVC, and $FEV_1$/FVC was calculated by taking the difference between precent predicted post-bronchodilator and pre-bronchodilator.
Abbreviations: $FEV_1$: Forced expiratory volume in 1 second; FVC: Forced vital capacity; TLC, Total lung capacity; $DL_{CO}$: Diffusing capacity for carbon monoxide

**Supplementary Table E4.** Distribution of oscillometry measurements in all individuals

| Oscillometry Measure | Mean (SD) | Median (25%, 75%) | Min, Max |
|---|---|---|---|
| **Pre-albuterol % predicted *** | | | |
| $R_5$ | 101.0 (30.4) | 100.8 (83.6, 118.0) | 48.9, 191.7 |
| $R_{19}$ | 98.5 (25.1) | 95.8 (80.5, 111.5) | 62.5, 165.6 |
| $X_5$ | 96.1 (91.8) | 76.3 (64.9, 94.7) | 37.9, 615.5 |
| AX | 164.3 (187.8) | 105.3 (74.8, 189.2) | 29.8, 1099.0 |
| $F_{res}$ | 105.8 (25.9) | 100.7 (83.7, 126.3) | 66.2, 164.4 |
| **Post-albuterol % predicted *** | | | |
| $R_5$ | 88.1 (22.1) | 84.9 (71.7, 105.8) | 47.0, 127.4 |
| $R_{19}$ | 87.2 (18.8) | 86.0 (74.5, 98.4) | 54.5, 135.5 |
| $X_5$ | 78.6 (52.5) | 67.8 (54.0, 87.9) | 25.9, 355.0 |
| AX | 116.4 (106.0) | 82.2 (55.0, 142.0) | 19.4, 595.6 |
| $F_{res}$ | 96.9 (21.5) | 92.2 (83.4, 112.3) | 61.8, 144.1 |
| **% change **** | | | |
| $R_5$ | -9.5 (22.4) | -13.1 (-28.1, 2.7) | -37.5, 46.7 |
| $R_{19}$ | -8.8 (19.4) | -11.4 (-24.3, -2.2) | -35.4, 39.5 |
| $R_5$ - $R_{19}$ | -42.7 (135.4) | -36.3 (-87.4, 41.9) | -550.0, 166.7 |
| $X_5$ | 11.0 (28.8) | 12.9 (-0.4, 29.4) | -87.3, 60.3 |
| AX | -14.8 (47.9) | -20.8 (-49.6, 8.3) | -75.6, 166.4 |
| $F_{res}$ | -6.4 (16.7) | -5.9 (-17.8, 2.1) | -44.0, 28.9 |

* Percent predicted and raw values are measurements taken before subject was given albuterol (pre-bronchodilator) except $R_5$-$R_{19}$.
** Percent change in value was calculated by taking the difference between precent predicted post-bronchodilator and pre-bronchodilator.
Abbreviations: $R_5$, Total airway resistance at 5 Hz; $R_{19}$, Central airway resistance at 19 Hz; $R_5$-$R_{19}$, Small (peripheral) airway resistance, frequency dependence of resistance; $F_{res}$, resonant frequency; AX, area of reactance; $X_5$, respiratory system reactance.

**Supplementary Table E5.** Percent predicted and percent change of oscillometry measurements in asymptomatic and persistent respiratory symptoms individuals

| | **Total n=40** | **Asymptomatic n=24** | **Symptomatic n=16** | |
|---|---|---|---|---|
| **Oscillometry Measure** | Mean (SD) Median (25%, 75%) | Mean (SD) Median (25%, 75%) | Mean (SD) Median (25%, 75%) | *p-value* * |
| **Pre-albuterol % predicted** ** | | | | |
| **$R_5$** | 101.0 (30.4) 100.8 (83.6, 118.0) | 93.7 (25.5) 92.0 (72.9, 114.4) | 112.1 (34.4) 107.4 (93.3, 118.5) | *0.09* |
| **$R_{19}$** | 98.5 (25.1) 95.8 (80.5, 111.5) | 94.8 (25.7) 93.2 (71.7, 106.0) | 104.0 (23.8) 100.6 (87.3, 117.6) | *0.18* |
| **$X_5$** | 96.1 (91.8) 76.3 (64.9, 94.7) | 76.3 (17.9) 75.6 (68.8, 79.9) | 125.8 (140.8) 76.7 (59.8, 125.7) | *0.63* |
| **AX** | 164.3 (187.8) 105.3 (74.8, 189.2) | 115.3 (75.1) 89.6 (73.7, 123.7) | 237.9 (270.9) 163.3 (80.1, 252.2) | *0.09* |
| **$F_{res}$** | 105.8 (25.9) 100.7 (83.7, 126.3) | 98.1 (22.1) 92.0 (81.7, 110.0) | 117.3 (27.5) 124.0 (94.0, 141.1) | ***0.01*** |
| **$R_5$ - $R_{19}$, cm $H_2O$/L/sec** ** | 0.4 (0.7) 0.3 (0.0, 0.5) | 0.2 (0.3) 0.1 (0.0, 0.4) | 0.7 (0.9) 0.4 (0.2, 0.7) | ***0.01*** |
| **Post-albuterol % predicted** ** | | | | |
| **$R_5$** | 88.1 (22.1) 84.9 (71.7, 105.8) | 83.9 (22.2) 79.4 (67.9, 104.1) | 94.2 (21.1) 88.1 (82.4, 113.4) | *0.16* |
| **$R_{19}$** | 87.2 (18.8) 86.0 (74.5, 98.4) | 86.9 (22.9) 79.7 (69.5, 104.2) | 87.8 (10.9) 89.2 (78.6, 96.5) | *0.49* |
| **$X_5$** | 78.6 (52.5) 67.8 (54.0, 87.9) | 66.4 (19.9) 65.9 (57.1, 75.1) | 96.9 (77.2) 83.4 (49.2, 114.5) | *0.16* |
| **AX** | 116.4 (106.0) 82.2 (55.0, 142.0) | 84.5 (43.3) 77.1 (55.0, 88.6) | 164.2 (149.3) 121.2 (54.5, 254.1) | *0.09* |
| **$F_{res}$** | 96.9 (21.5) 92.2 (83.4, 112.3) | 91.0 (14.5) 90.2 (81.7, 95.0) | 105.7 (27.3) 104.4 (88.2, 134.7) | *0.07* |
| **$R_5$ - $R_{19}$, cm $H_2O$/L/sec** *** | 0.3 (0.5) 0.1 (0.0, 0.5) | 0.1 (0.3) 0.1 (0.0, 0.3) | 0.6 (0.7) 0.4 (0.0, 0.8) | *0.06* |
| **% change** **** | | | | |
| **$R_5$** | -9.5 (22.4) -13.1 (-28.1, 2.7) | -7.6 (23.30) -12.1 (-27.7, 4.4) | -12.3 (21.50) -20.4 (-28.9, 1.1) | *0.56* |
| **$R_{19}$** | -8.8 (19.4) -11.4 (-24.3, -2.2) | -6.1 (21.4) -9.3 (-24.3, 1.5) | -12.8 (15.8) -13.4 (-27.9, -3.6) | *0.35* |
| **$R_5$ - $R_{19}$** | -42.7 (135.4) -36.3 (-87.4, 41.9) | -48.2 (162.4) -50.9 (-87.4, 85.3) | -34.5 (84.2) -34.5 (-106.3, 30.4) | *0.92* |
| **$X_5$** | 11.0 (28.8) 12.9 (-0.4, 29.4) | 11.2 (26.3) 12.0 (-3.3, 25.9) | 10.7 (33.2) 19.6 (6.1, 33.4) | *0.63* |
| **AX** | -14.8 (47.9) -20.8 (-49.6, 8.3) | -14.7 (42.2) -20.8 (-50.1,13.0) | -14.8 (56.9) -22.9 (-49.6, -4.1) | *0.67* |
| **$F_{res}$** | -6.4 (16.7) -5.9 (-17.8, 2.1) | -4.8 (17.4) -6.2 (-18.4, 4.1) | -8.9 (15.8) -5.9 (-17.8, -1.1) | *0.61* |

* Significant differences between asymptomatic and symptomatic groups determined by Mann-Whitney U test ($p < 0.05$).
** Percent predicted and raw values are measurements taken before subject was given albuterol (pre-bronchodilator).
*** This measurement was taken after subject was given albuterol (post-bronchodilator).
**** Percent change in value was calculated by taking the difference between precent predicted post-bronchodilator and pre-bronchodilator.
Abbreviations: $R_5$, Total airway resistance at 5 Hz; $R_{19}$, Central airway resistance at 19 Hz; $R_5$-$R_{19}$, Small (peripheral) airway resistance, frequency dependence of resistance; $F_{res}$, resonant frequency; AX, area of reactance; $X_5$, respiratory system reactance.

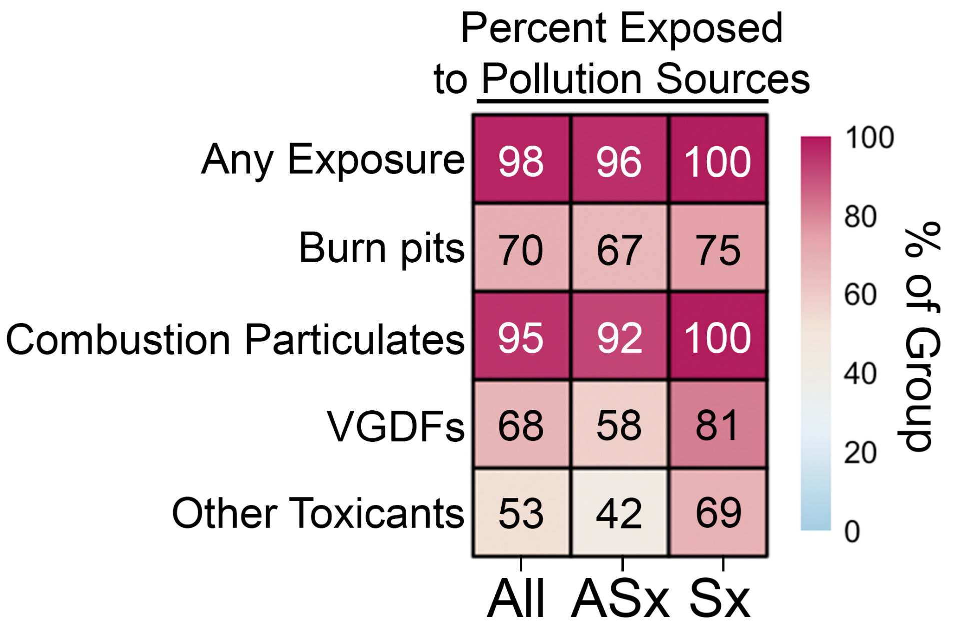


**Supplementary Figure E1. Percentage of individuals exposed to deployment-related airborne pollutant sources.** Forty non-smoker individuals with persistent (Sx, n=16) and without (ASx, n=24) respiratory symptoms participated in a standardized 32-item exposure battery assessment as part of the SHADE and LEDE studies. The exposure battery included questions related to sustained or direct exposure to burn pit smoke, open air combustion by-products other than burn pits (e.g., exposures related to combat, other military activities, oil well and refinery fires, or engine exhaust), military occupation-related vapors, gas, dusts, or fumes (VGDFs), or other toxicants (e.g., pesticides and chemical warfare agents). Percentage of individuals with affirmative responses to the above questions in the battery regarding exposure to different sources. Percentages in heatmap reflect median % within groups (e.g., % of all individuals, asymptomatics, and symptomatics). Significant differences between asymptomatic and symptomatic groups were determined by Mann-Whitney U test, *$p < 0.05$. No significant differences were detected between asymptomatic and symptomatic groups for percent exposure for each category.

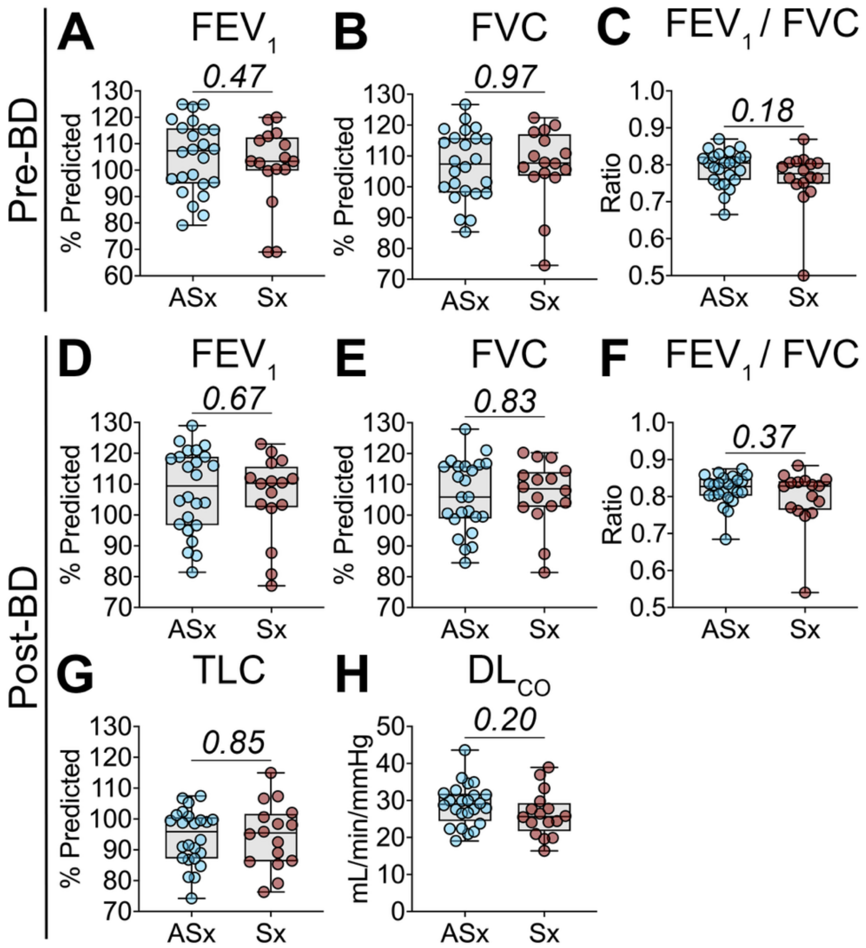


**Supplementary Figure E2. Pulmonary function tests assessed in pollutant-exposed asymptomatic and symptomatic individuals before and after bronchodilator (BD) administration.** Forty non-smoker individuals with persistent (Sx, n=16) and without (ASx, n=24) respiratory symptoms underwent pulmonary function tests as part of the SHADE and LEDE studies. PFTs were taken (**A-C**) before and (**D-H**) after administration of 180 µg of albuterol. PFT panel included: **(A, D)** Forced expiratory volume in 1 second ($FEV_1$), **(B, E)** forced vital capacity (FVC), **(C, F)** $FEV_1$/FVC ratio, **(G)** total lung capacity (TLC), and **(H)** diffusing capacity for carbon monoxide ($DL_{CO}$). Medians presented with minimum, maximum, and IQR. Significant differences between asymptomatic and symptomatic groups were determined by unpaired Student's t-test, *$p < 0.05$.

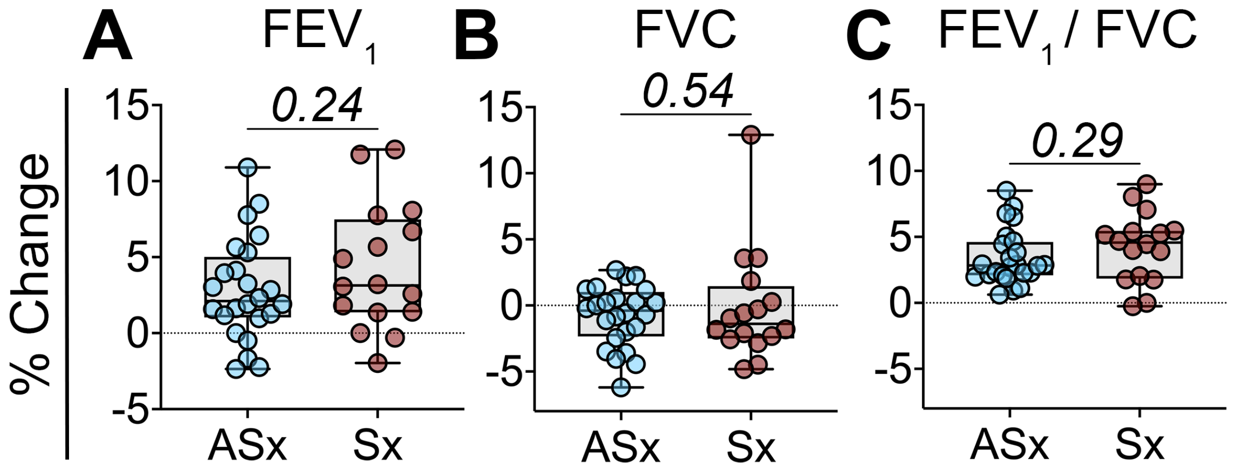


**Supplementary Figure E3. Percent change of each pulmonary function tests between pre- and post-albuterol administration in pollutant-exposed asymptomatic and symptomatic individuals.** Forty non-smoker individuals with persistent (Sx, n=16) and without (ASx, n=24) respiratory symptoms underwent pulmonary function tests. Percent change in value for **(A)** $FEV_1$, **(B)** FVC, and **(C)** $FEV_1$/FVC ratio was calculated by taking the difference between precent predicted of PFTs taken before and after administration of 180 µg of albuterol for each subject. Medians presented with minimum, maximum, and IQR. Significant differences between asymptomatic and symptomatic groups were determined by unpaired Student t-test, *$p < 0.05$.

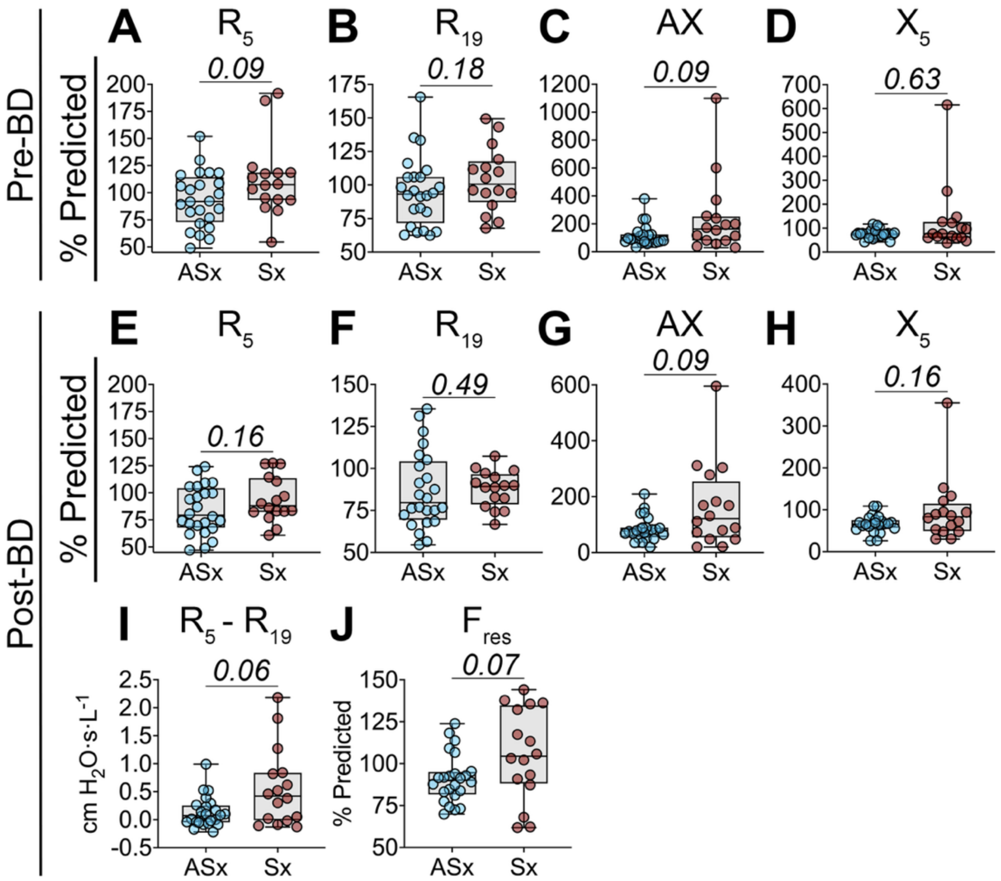


**Supplemental Figure E4. Distribution of oscillometry measurements in asymptomatic and symptomatic individuals.** Forty non-smoker individuals with persistent (Sx, n=16) and without (ASx, n=24) respiratory symptoms underwent oscillometry measurements as part of the SHADE and LEDE studies. Measurements were taken (**A-D**) before and (**E-J**) after bronchodilator (BD) administration of 180 μg of albuterol. Panel included: **(A, E)** total airway resistance at 5 Hz ($R_5$), **(B, F)** central airway resistance at 19 Hz ($R_{19}$), **(C, G)** area of reactance (AX), **(D, H)** reactance at 5 Hz ($X_5$), **(I)** frequency dependence of resistance ($R_5$-$R_{19}$), and **(J)** resonant frequency ($F_{res}$). Medians are presented with minimum, maximum, and IQR. Significant differences between asymptomatic and symptomatic groups were determined by Mann-Whitney U test, *$p < 0.05$.

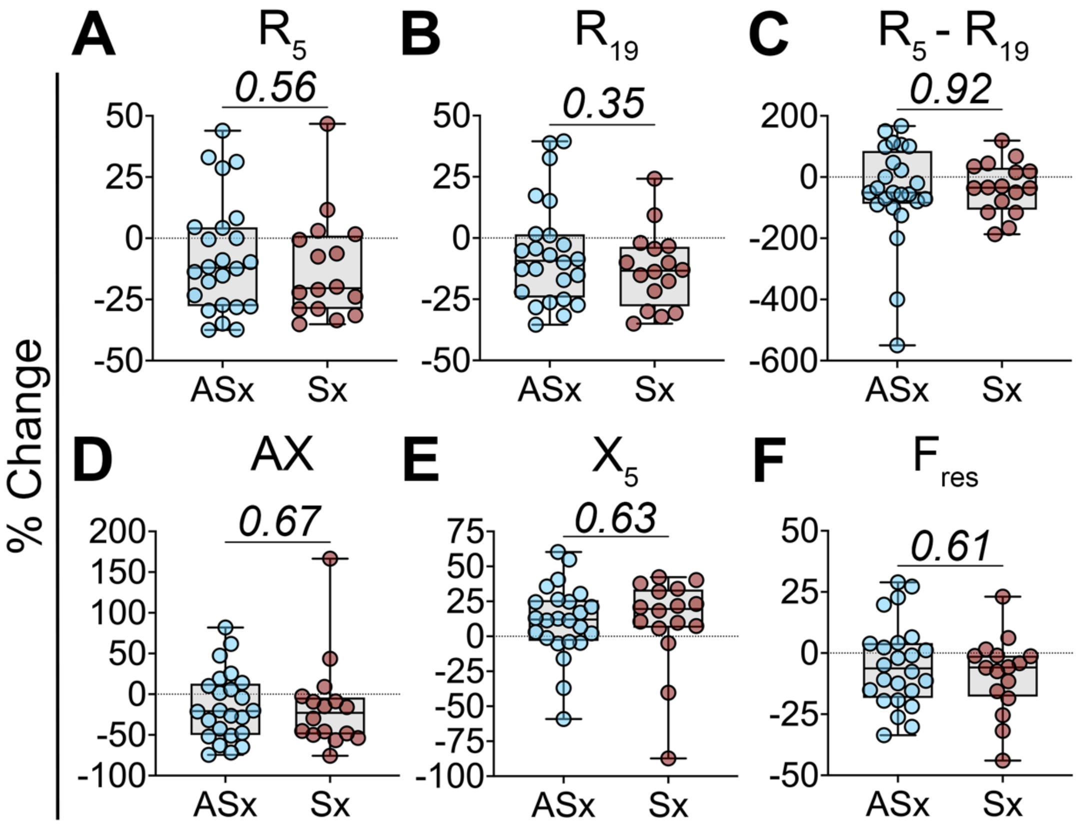


**Supplementary Figure E5. Percent change of each oscillometry measurement between pre- and post-albuterol administration in pollutant-exposed asymptomatic and symptomatic individuals.** Forty non-smoker individuals with persistent (Sx, n=16) and without (ASx, n=24) respiratory symptoms underwent oscillometry measurements. Percent change in value for **(A)** total airway resistance at 5 Hz ($R_5$), **(B)** central airway resistance at 19 Hz ($R_{19}$), **(C)** frequency dependence of resistance ($R_5$-$R_{19}$), (**D**) area of reactance (AX), (**E**) reactance at 5 Hz ($X_5$), and (**F**) resonant frequency ($F_{res}$) was calculated by taking the difference between precent predicted of PFTs taken before and after administration of 180 µg of albuterol for each subject. Medians are presented with minimum, maximum, and IQR. Significant differences between asymptomatic and symptomatic groups were determined by Mann-Whitney U test, **p* < *0.05*.

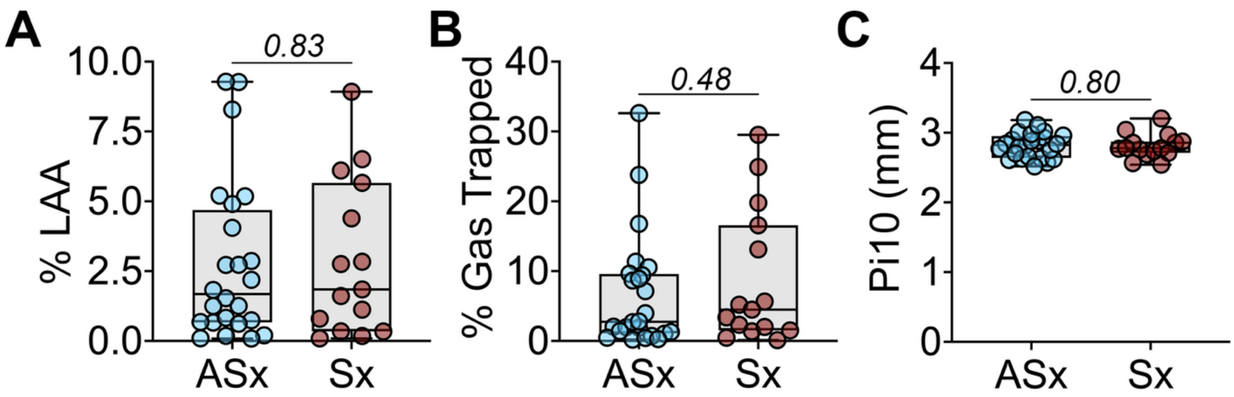


**Supplemental Figure E6. Computed tomography and airway thickness assessment exhibited no differences in symptomatic compared to asymptomatic group.** Computed tomography was performed on forty individuals with persistent (Sx, n=16) and without (ASx, n=24) respiratory symptoms to measure **(A)** low attenuation area (% LAA), **(B)** gas trapped percentage, and **(C)** airway thickness (Pi10). Medians are presented with minimum, maximum, and IQR. Significant differences between asymptomatics and symptomatic groups were determined by Mann-Whitney U test, $^*p < 0.05$.

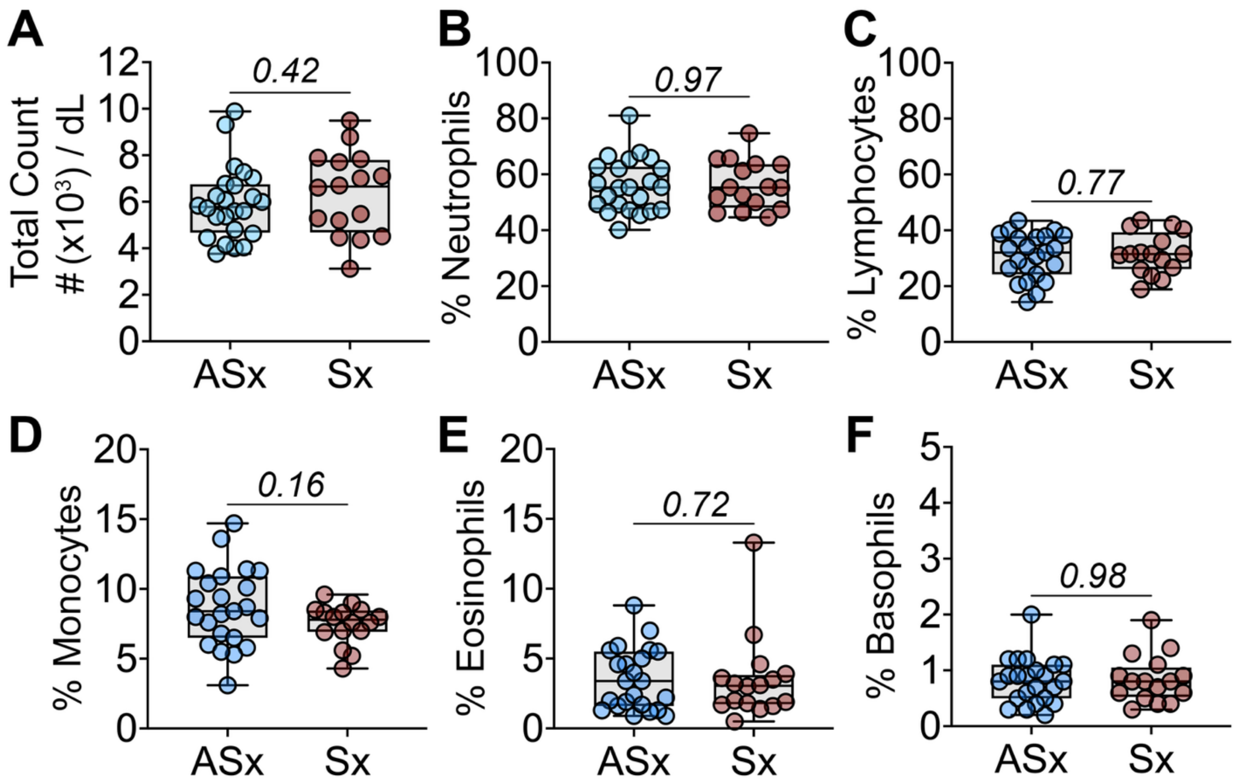


**Supplemental Figure E7. Neutrophils and lymphocytes comprise majority of white blood cells in pollutant-exposed individuals.** WBC differential count was performed on blood isolated from asymptomatic (ASx, n=24) and symptomatic (Sx, n=16) individuals to measure **(A)** total white blood cells, **(B)** neutrophils, **(C)** lymphocytes, **(D)** monocytes, **(E)** eosinophils, and **(F)** basophils. Medians are presented with minimum, maximum, and IQR. Significant differences between asymptomatic and symptomatic groups were determined by Mann-Whitney U test, $^*p < 0.05$.

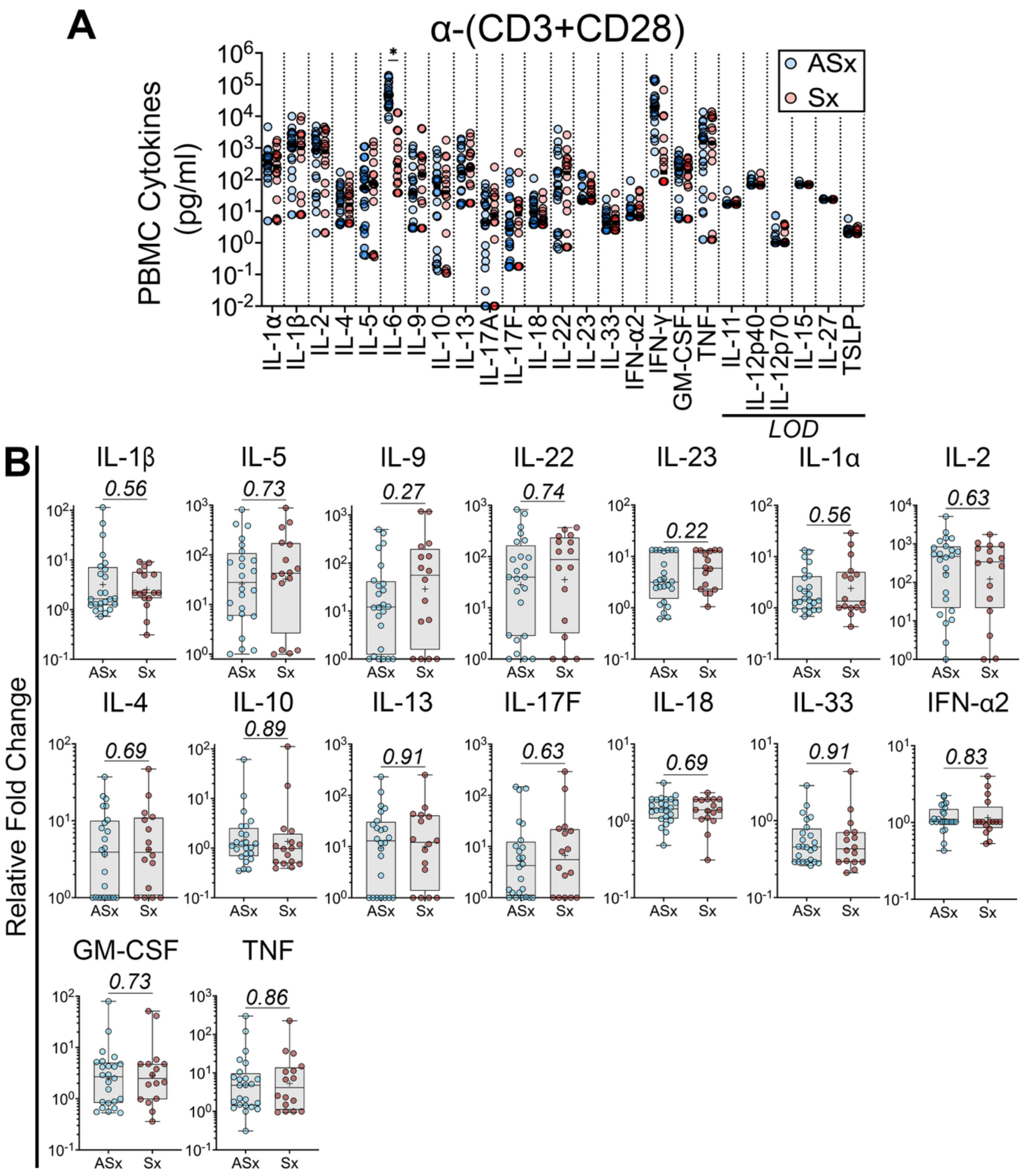


**Supplemental Figure E8. Adaptive immune activation of PBMCs from pollutant-exposed individuals.** *Ex vivo* immune activation was assessed using PBMCs ($10^6$ cells/well in triplicates) from asymptomatic (ASx, n=24) and symptomatic (Sx, n=16) individuals. PBMCs were stimulated with T-lymphocyte coactivators (α-CD3 and α-CD28 antibodies, 1 μg/ml) for 36 hours before supernatant collection. **(A)** Cytokine levels (pg/ml)

of each individual were then measured using the LegendPlex multiplex kit. Horizontal line represents median. **(B)** Individual data points of log10-transformed relative fold changes are displayed for cytokines exceeding limit of detection (LOD). Boxplot displays median with minimum, maximum, and IQR and + represents mean. Fold change was normalized to each individual's unstimulated PBMCs (baseline) before log10-transformation. For **A-B**, unpaired Student's t-test was used to determine significant differences between the two groups, *$p < 0.05$.

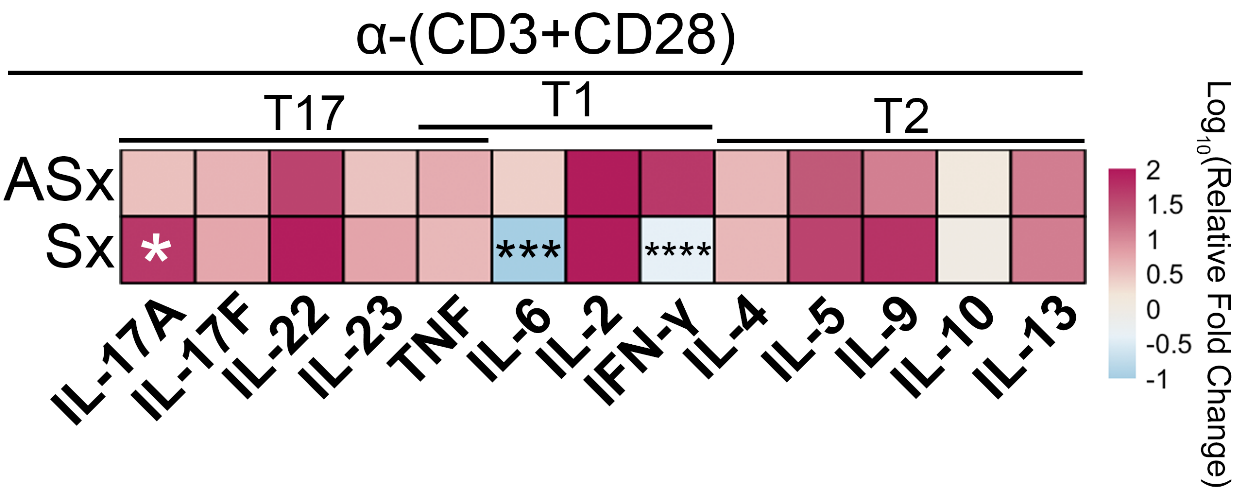


**Supplemental Figure E9. Symptomatic individuals exhibited a mixed T2 and T17 with reduced T1 phenotype.** PBMCs were stimulated with T-lymphocyte coactivators (α-CD3 and α-CD28 antibodies, 1 μg/ml) for 36 hours before supernatant collection. Cytokine levels were then measured using the LegendPlex multiplex kit. Heatmaps display cytokines associated with type 1 (T1), type 2 (T2), and type 17 (T17) signatures representing $\log_{10}$ of medians in relative fold changes from asymptomatic (n=24) and symptomatic (n=16) individuals. Cytokines exceeding threshold for limit of detection are displayed. Fold change was normalized to each individual's unstimulated PBMCs (baseline) before log10-transformation. Unpaired Student's t-test was used to determine significant differences between the two groups, **p < 0.05*, ***p<0.01,* ****p<0.001*, *****p<0.0001.*

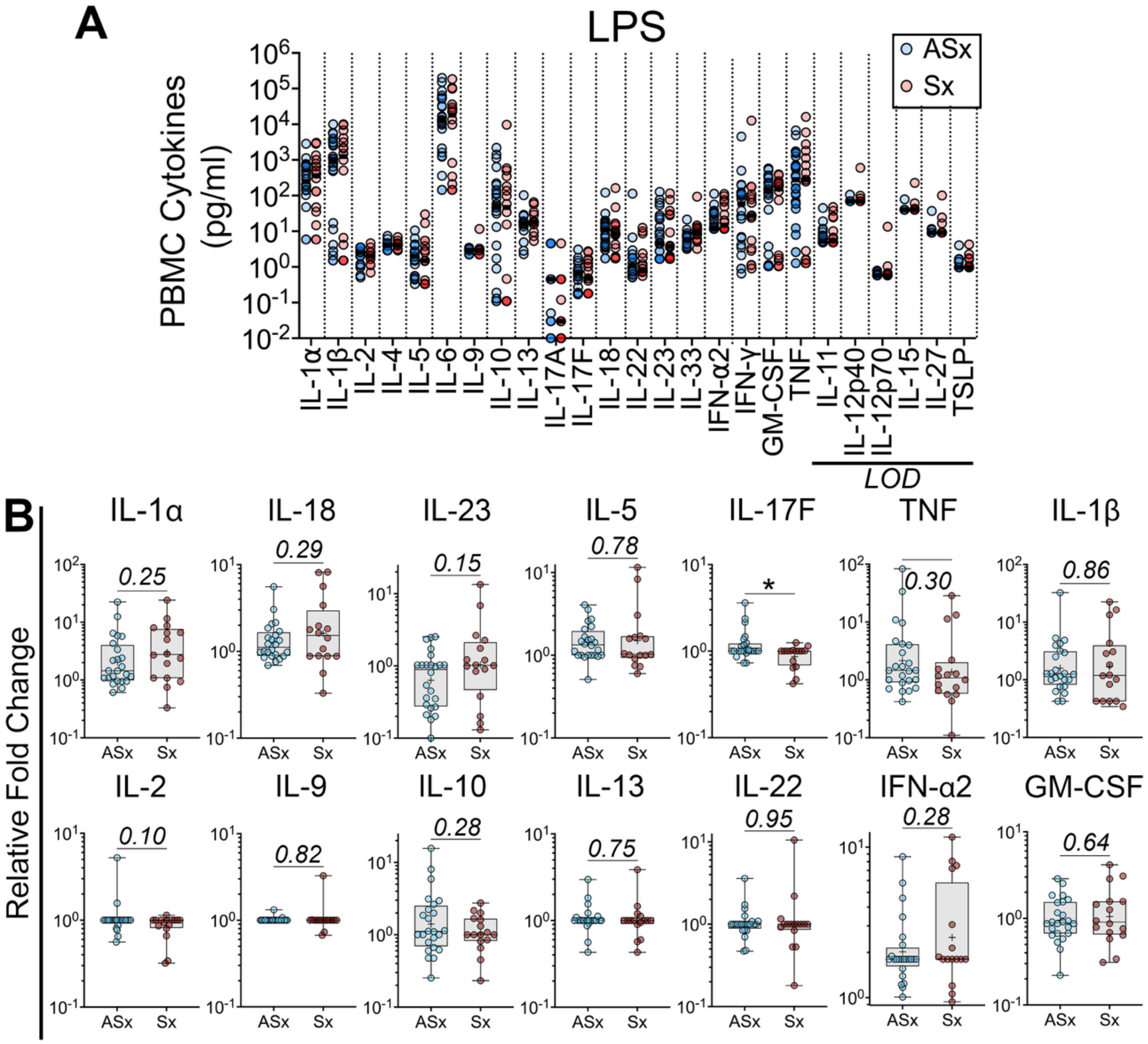


**Supplemental Figure E10. PBMCs from pollutant-exposed individuals were stimulated with LPS.** *Ex vivo* immune activation was assessed using PBMCs ($10^6$ cells/well in triplicates) from asymptomatic (ASx, n=24) and symptomatic (Sx, n=16) individuals. PBMCs were stimulated with LPS (100 ng/ml) for 36 hours before supernatant collection. **(A)** Cytokine levels (pg/ml) of each individual were then measured using the LegendPlex multiplex kit. Horizontal line represents median. **(B)** Individual data points of log10-transformed relative fold changes are displayed for cytokines exceeding limit of detection (LOD). Boxplot displays median with minimum, maximum, and IQR and +

represents mean. Fold change was normalized to each individual's unstimulated PBMCs (baseline) before log10-transformation. For **A-B**, unpaired Student's t-test was used to determine significant differences between the two groups, *$p < 0.05$.

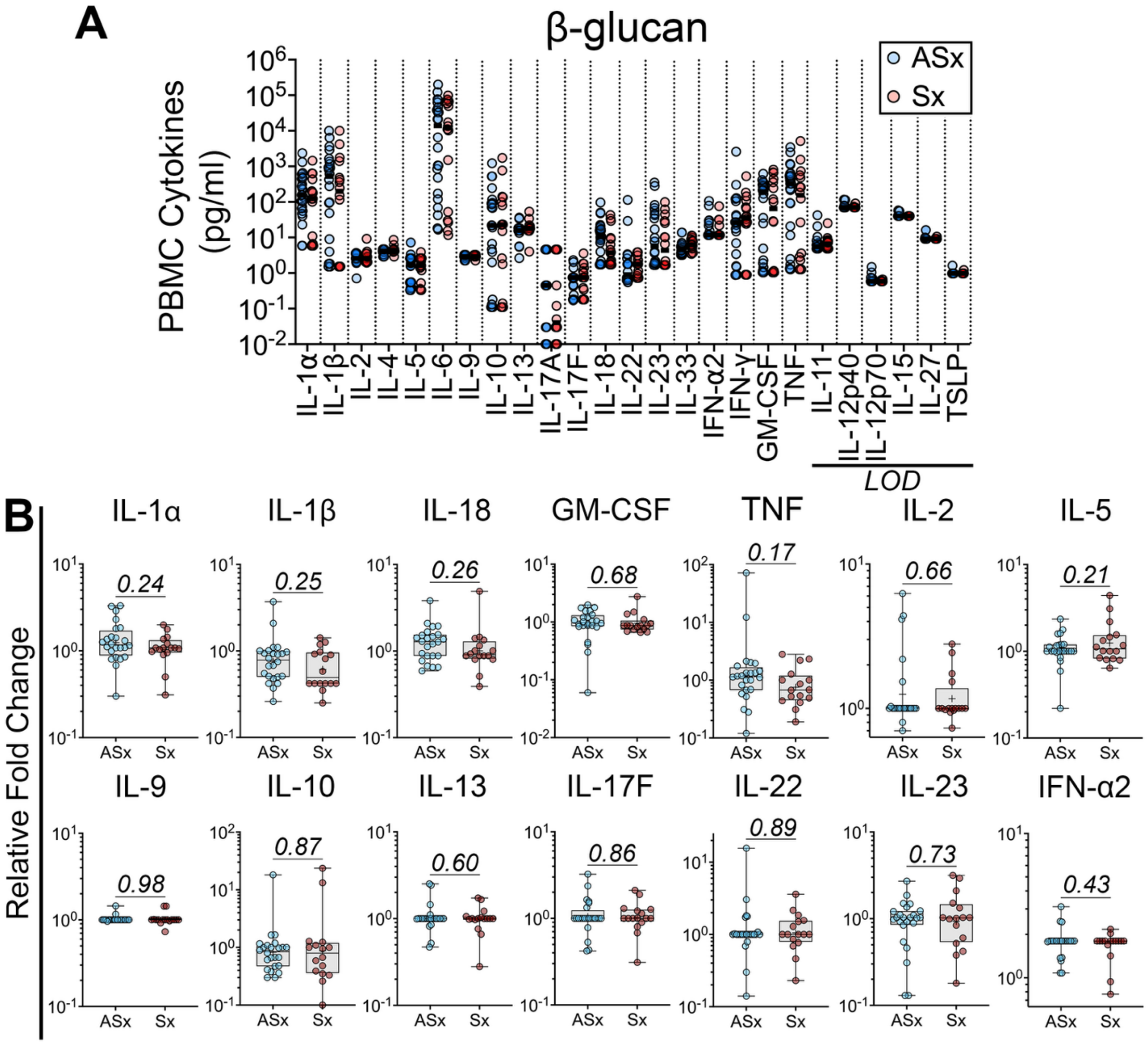


**Supplemental Figure E11. PBMCs from pollutant-exposed individuals were stimulated with β-glucan.** *Ex vivo* immune activation was assessed using PBMCs ($10^6$ cells/well in triplicates) from asymptomatic (ASx, n=24) and symptomatic (Sx, n=16) individuals. PBMCs were stimulated with β-glucan (1 µg/ml) for 36 hours before supernatant collection. **(A)** Cytokine levels (pg/ml) of each individual were then measured using the LegendPlex multiplex kit. Horizontal line represents median. **(B)** Individual data points of log10-transformed relative fold changes are displayed for cytokines exceeding limit of detection (LOD). Boxplot displays median with minimum, maximum, and IQR and

+ represents mean. Fold change was normalized to each individual's unstimulated PBMCs (baseline) before log10-transformation. For **A-B**, unpaired Student's t-test was used to determine significant differences between the two groups, $*p < 0.05$.

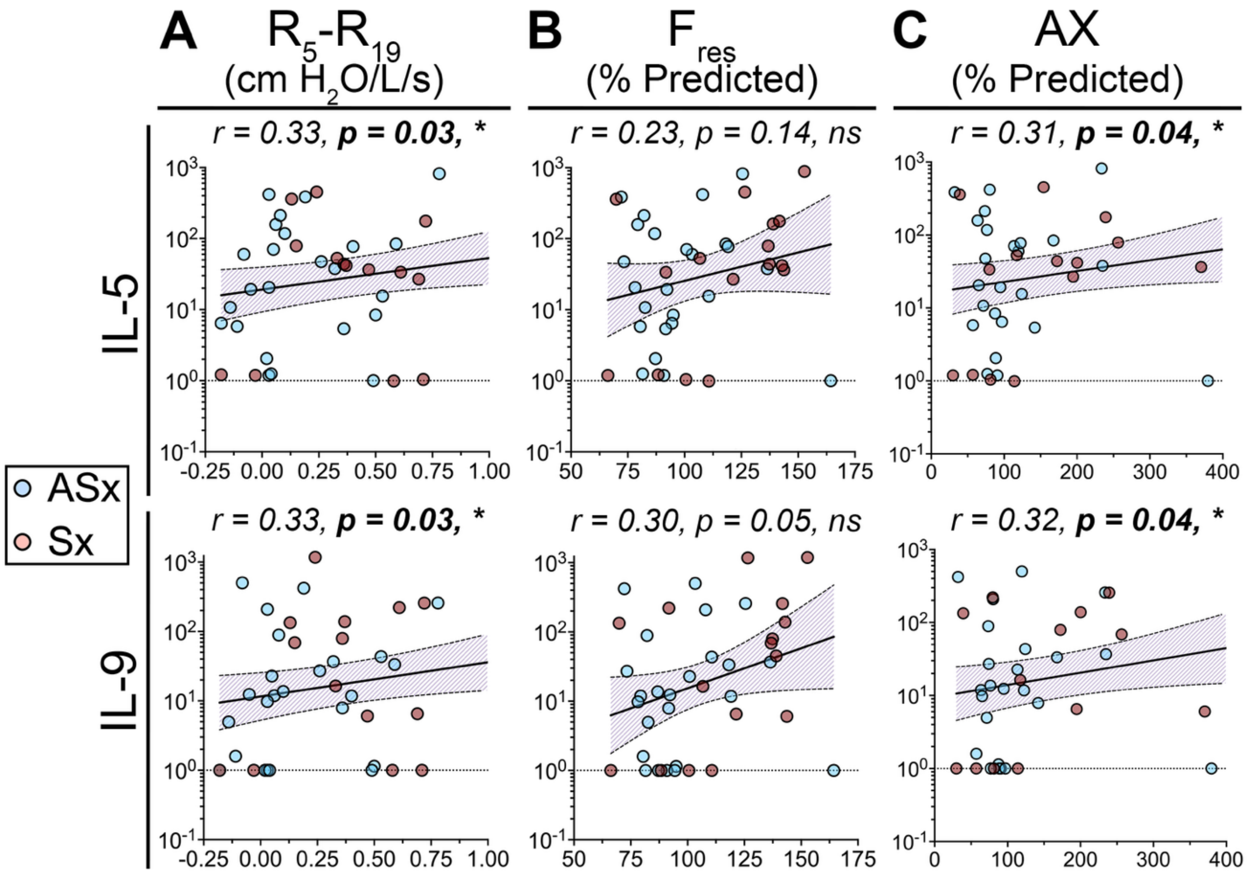


**Supplemental Figure E12. T2-associated cytokines positively correlated with increased small airway resistance and decreased compliance in pollutant-exposed individuals.** Correlation analysis was performed in pollutant-exposed individuals to determine correlation strength (*r*) between $log_{10}$-fold change of T2-associated cytokines IL-5 and IL-9 in α-(CD3+CD28) antibodies with pre-bronchodilator (**A**) small airway resistance ($R_5$-$R_{19}$), (**B**) resonant frequence ($F_{res}$), and (**C**) area of reactance (AX). Each data point represents each asymptomatic (n = 24, blue dots) or symptomatic (n = 16, red dots) individual. Dashed line represents linear fold change of 1, which is unstimulated PBMCs. The shaded region represents error bands within 95% confidence interval. Pearson correlation analysis was performed to determine significance. *$p < 0.05$.*
